# Promiscuous RNA binding by WDR5 remodels the KMT2A (MLL1) histone methyltransferase complex to an inactive state

**DOI:** 10.64898/2026.05.22.727156

**Authors:** Amoldeep S. Kainth, Pallavi Sirjoosingh, Michael S. Werner, Ankit Gupta, Akiko Koide, Shohei Koide, Alexander J. Ruthenburg

## Abstract

Chromatin-modifying complexes are critical in gene regulation, yet their proposed interactions with RNA remain poorly understood. Here, we show that WDR5, an essential subunit of the MLL1 (KMT2A) histone methyltransferase complex, binds RNA with high affinity but without sequence specificity. Using a stringent approach, we demonstrate that WDR5 directly engages a diverse pool of RNAs in cells, predominantly as a function of RNA abundance rather than specific motifs. Equilibrium binding assays further show that RNA length, rather than sequence, dictates WDR5 affinity. We map multiple RNA-binding regions on WDR5, some of which overlap with surfaces used in MLL1 complex subunit interfaces. Strikingly, we find that RNA binding disrupts the MLL1 complex by competitively displacing WDR5 from these critical protein-protein interactions, leading to a marked inhibition of MLL1 catalytic activity. This regulatory mechanism provides a layer of control over histone methylation, potentially integrating transcriptional activity with chromatin state.

## INTRODUCTION

Chromatin modifying complexes play crucial roles in gene regulation, development, and disease by altering the structure and accessibility of chromatin^1,2^, yet the mode of recruitment and regulation of many of these complexes at the target genomic loci remains an active area of research. Recent studies suggest that RNA, especially the non-coding RNA, interact with many chromatin-modifying complexes and may serve as potential regulators by influencing their activity and localization^3–8^. Therefore, a deeper understanding of the molecular underpinnings of these interactions – such as sequence specificity, interaction surfaces, and functional outcomes – could not only provide valuable insights into the regulation of chromatin modifying complexes but also present better therapeutic targets than the conventionally targeted sites which have constantly encountered clinical resistance^9–12^.

Among the chromatin-modifying complexes, the MLL (also known as SET1/KMT2 or COMPASS) family of histone methyltransferases (HMTs) is of particular interest due to its pivotal role in transcriptional activation by catalyzing the methylation of lysine 4 on histone H3 (H3K4me) ^13–15^. The mammalian MLL family comprises six members – MLL1 (KMT2A), MLL2 (KMT2B), MLL3 (KMT2C), MLL4 (KMT2D), SET1A (KMT2F), and SET1B (KMT2G) – that share a common set of core subunits: WDR5, RBBP5, ASH2L, and DPY30^16,17^. Biochemical studies have shown that these core components are essential for methyltransferase activity across MLL complexes, albeit to varying degrees – RBBP5 and ASH2L are required for all MLL complexes, while WDR5 is specifically essential for MLL1^18–26^. MLL1 is shown to play a key role in differentiation^27^, development^28–31^ and disease progression^32–35^. Multiple mechanisms have been proposed for the recruitment of the MLL1 complex to its genomic targets, including interactions with Menin^21,36^, specific recognition domains within the MLL1 polypeptide^37–41^, and direct RNA binding by the WDR5 subunit^42–48^.

As the formative example of WDR5-RNA mediated recruitment of MLL1 complex, the long noncoding RNA (lncRNA) HOTTIP is proposed to be a *cis-*regulator of the distal portion of the *HOXA* gene cluster^42,44,46^. While RNA depletion studies have established that HOTTIP and other lncRNAs are essential for recruiting the MLL1 complex^42,44,46,47,49,50^, a clear demonstration that such a recruitment is mediated by sequence-specific interaction between RNA and WDR5 subunit of the MLL1 complex is lacking. Prior studies claiming specificity notably lack equilibrium binding assays with appropriate negative controls on both the RNA and protein side of the interface^42,44^. Consistent with the lack of specificity, no consensus RNA motif is discovered despite numerous attempts. Therefore, rigorous biochemical experiments are required to determine the strength, specificity and functional implications of RNA binding to WDR5.

The need for robust molecular understanding of RNA-WDR5 interactions is clearly demonstrated by recent investigations and debate on the validity, extent and functional significance of the interaction between RNA and chromatin regulators. A key point of disagreement is the strength of evidence for direct binding and functional significance of protein-RNA interactions in cells for proteins shown to bind RNA *in vitro* exemplified by the PRC2 complex^51–55^. Early reports suggested that PRC2 is recruited by specific binding to RepA motif of *Xist* RNA to deposit the repressive H3K27me3 marks at the silenced X chromosome^3^. Whereas several studies claimed that PRC2 binds promiscuously to a wide range of RNAs with little to no sequence specificity^56–58^. More recent investigations show that subtle biochemical staging differences can account for some of these discrepancies^51^, or argue that G-rich RNA sequences, particularly those which can form RNA G-quads are involved in the recruitment^59–61^. Challenging these models altogether, other cell-based studies relegate the detection of PRC2-RNA interactions to be completely artifactual^62^ or indirect^63,64^. The discrepancies between the cell-based studies can be attributed to biases in the crosslinking efficiencies of the crosslinkers, biochemical artifacts of genomic approaches, use of arbitrary thresholds to demarcate actual binding from background RNA-protein interactions, and inherent limitations of characterizing non sequence-specific yet potentially functional binding. The contradictory PRC2 literature highlight the challenges of detecting *bona fide* RNA-protein interactions using stringent immunoprecipitation conditions, employing crosslinker independent methods, testing strength and specificity of binding by equilibrium quantitative binding measurements, and structure-guided mutagenesis to map the location and function of RNA binding to the protein. By combining each of these approaches, we sought to gain a more thorough understanding of WDR5-RNA binding and the functional consequences of this interaction for SET/MLL family histone methyltransferase activities.

Employing a stringent variant of PAR-CLIP, we detected a wide range of RNAs that crosslink directly to WDR5, generally in proportion to their nuclear abundance. Importantly, using a crosslink independent *in situ* method, we found that RNA is necessary for chromatin binding of WDR5. Furthermore, using quantitative equilibrium measurements we show that the *in vitro* binding affinity of WDR5 to model RNA resides in a biologically consequential mid-nanomolar range. Remarkably, this tight binding lacks any detectable sequence or structural contributions to apparent specificity, rather the length of an RNA-binding partner is the sole predictor of affinity. Structure-guided fine-grained mapping delineates multiple RNA-binding interfaces on WDR5, some of which overlap with binding surfaces of other subunits of the MLL1 complex, resulting in competitive binding to WDR5. Consequentially, RNA binding by WDR5 competitively inhibits the assembly of the catalytically competent MLL1 core HMTase complex causing a steep erosion of MLL1 enzymatic activity. Taken together, our results suggest markedly different role of RNA in chromatin regulation by which WDR5-RNA binding interferes with the MLL1-WDR5 binding interface, destabilizing the MLL1 complex, and thereby regulates MLL1 histone methyltransferase activity.

## RESULTS

### WDR5 directly binds RNA in cells with limited specificity

Prior experiments attempting to understand the nature and function of WDR5 interaction with RNA used formaldehyde crosslinking procedures^4,44^. However, these approaches were recently argued to lack sufficient stringency for unambiguous interpretation, as they cannot distinguish direct from indirect binding of WDR5 to RNA via another RNA-binding subunit in the same protein complex^62^ or spurious RNA-RNA crosslinking by formaldehyde^65^. To map the pool of RNA that directly engages WDR5 in living cells, we developed PAR-CRAC which is a hybrid of PAR-CLIP (*<u>P</u>*hoto-*<u>A</u>*ctivatable *<u>R</u>*ibonucleoside-enhanced *<u>C</u>*ross*<u>L</u>*inking and *I*mmuno*<u>P</u>*recipitation) ^66^ and CRAC (*<u>Cr</u>*oss-linking *<u>a</u>*nalysis of *<u>c</u>*DNAs) ^67^, to cross-link and isolate WDR5-bound RNA species. Specifically, we used UV-mediated crosslinking to 4-thiouridine, with stringent tandem chromatography-based isolation of complexes under native, then denaturing conditions with FLAG-His_6_ tagged WDR5 stably expressed in HEK293 cells (Figure 1A, S1A). The crosslinked RNAs were then subjected to cDNA synthesis and high throughput sequencing. We note that a similar method, denaturing CLAP (*C*ovalent *L*inkage and *A*ffinity *P*urification), recently measured WDR5-RNA interaction in cells. We observed an overall good correlation between WDR5 PAR-CRAC and CLAP (Figure S1B). Based on read counts, we found that WDR5 binds to a large repertoire of RNA in cells extending well beyond lncRNAs, as previously suggested^4^ (Figure 1B-C, S1C). In addition to read counts, we quantified T > C conversion resulting from UV-crosslinking of thiouridine to protein^66,68^, to probe for WDR5-RNA interaction with higher stringency and resolution, independent of variations in the input sample (Figure 1B, S1D). When probing for the primary determinant of WDR5’s apparent binding to a given RNA, we observed that WDR5 PAR-CRAC signal was most correlated with the total RNA abundance (Figure 1C). Notably, we observed that the total amount of RNA was a better predictor of RNA capture than the gene-length normalized abundance or length of RNA, indicating that RNA binding to WDR5 is driven by bulk abundance. We did not find any evidence of an enriched sequence motif or potential RNA secondary structure such as RNA G-quadruplex that could account for WDR5-RNA interaction (Figure S1E, F). Similar to PAR-CRAC, signal counts of WDR5 denaturing CLAP showed highest correlation with RNA abundance (Figure S1G). Collectively these data are incongruent with models^42,44,46^ of RNA-mediated locus-specific recruitment of WDR5 that are inherently premised on the sequence-specific interaction of WDR5 and RNA.

**Figure 1.**
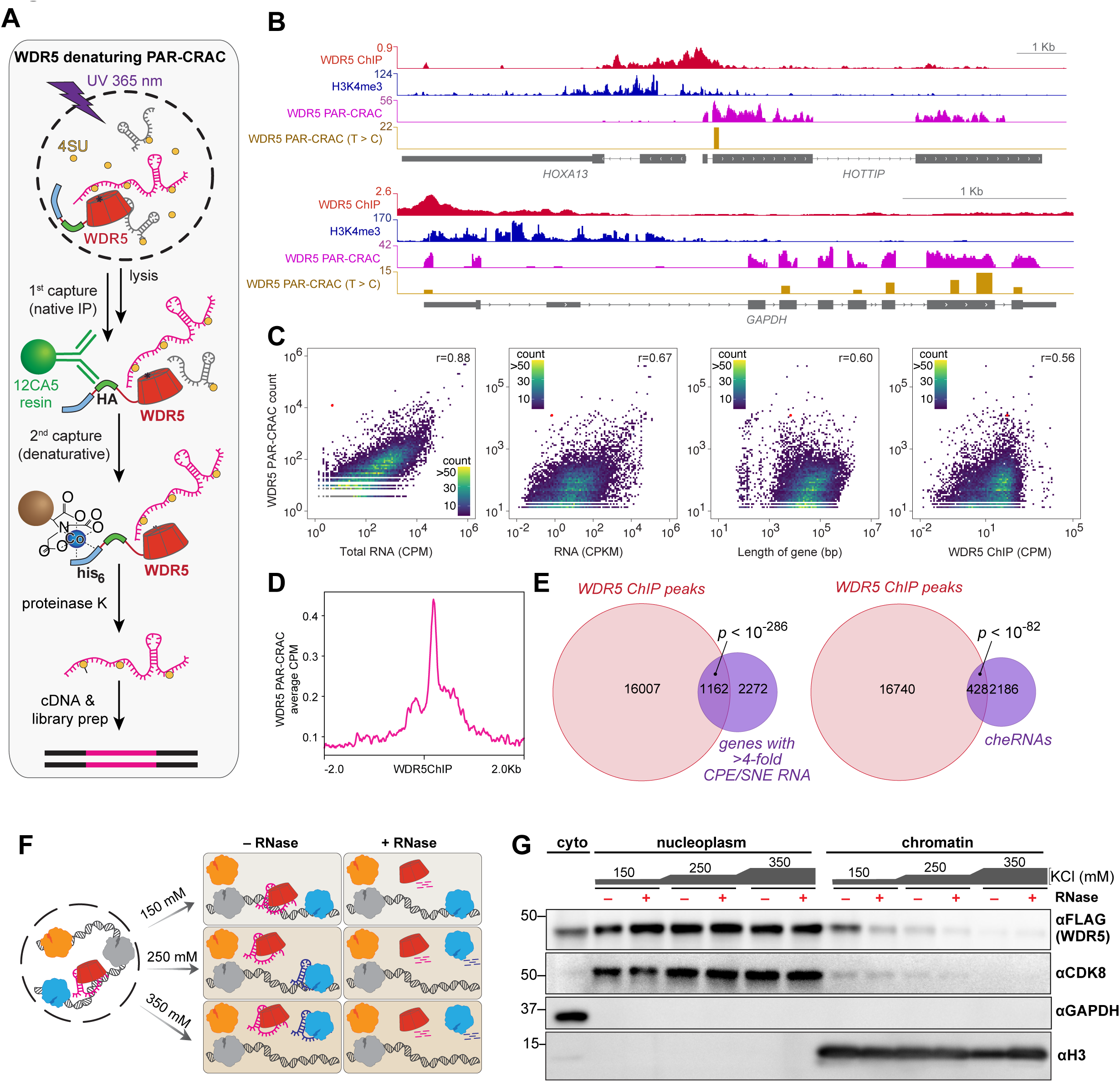
Promiscuous RNA binding of WDR5 in cells plays a role in its chromatin engagement. **(A)** Schematic of WDR5 denaturing PAR-CRAC (*<u>P</u>*hoto-*<u>A</u>*ctivatable *<u>R</u>*ibonucleoside-enhanced *<u>C</u>R*ossIinking and *<u>A</u>*nalysis of *<u>c</u>*DNAs) to stringently define cellular RNA-binding partners. 4-thiouridine (4SU), incorporated in the RNA of growing cells, was subjected to protein crosslinking at 365 nm. Epitope tagged WDR5 was then immunoprecipitated, first under native conditions using 12CA5 antibody (for the HA tag) and then under denaturing conditions (7.3 M Urea, 0.9M NaCl, 50°C) using Cobalt (Co^2+^) beads (for the 6X histidine tag). WDR5-bound RNA was then purified and subjected to high throughput sequencing. **(B)** View of the locus of origin of WDR5-bound RNA showing total count (PAR-CRAC) and crosslink-mediated thymine to cytosine count (T > C) at the *HOTTIP-HOXA13* (*top*) and the *GAPDH* (*bottom*) loci. WDR5 and H3K4me3 occupancy are shown for reference. **(C)** Correlation between WDR5 PAR-CRAC signal and indicated parameter of genes. CPM, counts per million mapped reads; CPKM, counts per thousand base pair of gene per million mapped reads. Only the genes with non-zero count in the comparison dataset are shown. Colors correspond to kernel density estimations of scatter plot distribution. r, Spearman correlation coefficient. **(D)** Average count of WDR5 PAR-CRAC count centered on WDR5 ChIP peaks. **^(E)^** Overlap of WDR5 ChIP peaks and (*left*) genomic regions with >4 fold enrichment of counts in the chromatin pellet relative to the soluble nuclear extract; (*right*) genomic regions emanating chromatin enriched RNA (cheRNA). ^69^ **(F)** Schematic of nuclear extraction under a range of salt concentrations coupled to RNase (A, I, T, and H) treatment. **(G)** Representative western blot showing the presence of indicated proteins in the cytoplasm (cyto), nucleoplasm and chromatin fraction after fractionation with the indicated salt concentrations in the absence or presence of RNase (A, I, T, and H) treatment. See also Figure S1

Next, we compared PAR-CRAC signal with the abundance of RNA in either the nucleoplasmic or chromatin bound pools derived from sub-nuclear biochemical fractionation of RNA^69^. As both chromatin-enriched and nucleoplasmic RNAs efficiently crosslinked with WDR5 in rough proportion to the abundance of the RNA species in each nuclear sub-compartment (Figure S1H), we infer that WDR5 likely engages both the chromatin bound and the nucleoplasmic pools of RNA in the nucleus. Despite substantial crosslinking of WDR5 to a large cohort of RNA as an approximate function of abundance, a small set of RNAs display marked enrichment in the WDR5 crosslinked pool, perhaps hinting at some specificity in binding (Figure S1H, S1I). Notably, the lncRNA HOTTIP is among the most overrepresented WDR5-crosslinked RNA versus any comparison group (ranging from 450-500-fold), consistent with the previous report that HOTTIP recruits the MLL1 complex to the distal portion of the *HOXA* cluster via interactions with WDR5^42,44^.

While the functional significance of WDR5-RNA interactions remains controversial^42,44,62^, several lines of data support the interpretation that WDR5 binds RNA in living cells, especially at the chromatin interface. First, there is a substantial overrepresentation of called WDR5 ChIP-seq peaks in HEK293 cells at gene loci that produce WDR5-crosslinked RNA (Figures 1D) and genes whose RNA is chromatin enriched (Figure 1E). Further, RNA emanating from WDR5-bound regions evinced higher enrichment in the chromatin fraction than in the nucleoplasmic fraction (Figure S1J). To directly test the role of RNA in the chromatin occupancy of WDR5, we performed nuclear fractionation with increasing salt concentration coupled to RNase treatment (Figure 1F). We observed that addition of RNase led to a reduction of WDR5 in the chromatin fraction with a corresponding increase in the nucleoplasm fraction, particularly at physiological salt regimes (Figure 1G, S1K), confirming that RNA plays a critical role in the chromatin occupancy of WDR5. RNase-dependent solubilization of a U2 spliceosomal component (SF3B155) from chromatin^70^ (Figure S1K) and no RNase-dependent change in histone 3 or mediator kinase (CDK8, Figure 1G) affirm the efficacy of this assay.

Taken together, our data suggest that WDR5 binds to a wide range of RNA in nuclei in approximate proportion to their abundance, and that RNA can contribute to the chromatin occupancy of WDR5, which not only resolves the conundrum of whether or not WDR5 binds to RNA in cells but also hints at the biological relevance of WDR5-RNA interaction.

### WDR5-RNA binding is promiscuous *in vitro*

Despite the abundance being the primary driver of WDR5-RNA interaction, HOTTIP emerged as a key outlier in PAR-CRAC dataset with disproportionate enrichment (Figure S1I). Owing to this enrichment and previously reported regulation of distal *HOXA* gene cluster transcription^42,44–46,48,71,72^, we chose HOTTIP as the model system to systematically explore the molecular determinants of affinity and specificity *in vitro*. First, we performed quantitative equilibrium filter binding assays with run-off transcribed HOTTIP RNA and fragments thereof (Figure 2A). We adopted forcing conditions to ensure the purified WDR5 was nucleic acid-free (see Methods) and confirmed that the complete MLL1 core histone methyltransferase complex (MLL1 [3745-3969], Ash2L, RbBP5, and Dpy30) ^24,73,74^ with our purified WDR5 is enzymatically active on histone 3 peptide (Figure S2A). Previous qualitative studies indicate that among the various subunits of MLL1 complex^42^, WDR5 accounts for the preponderance of RNA-binding. In our *in vitro* quantitative analysis of the RNA-binding comparison between WDR5 alone and the complete MLL1 complex, we find slightly tighter RNA-binding by the MLL1 complex as compared to WDR5 alone (Figure S2B) suggesting that WDR5-RNA binding represents the majority of the RNA-binding properties of the MLL1 complex.

**Figure 2.**
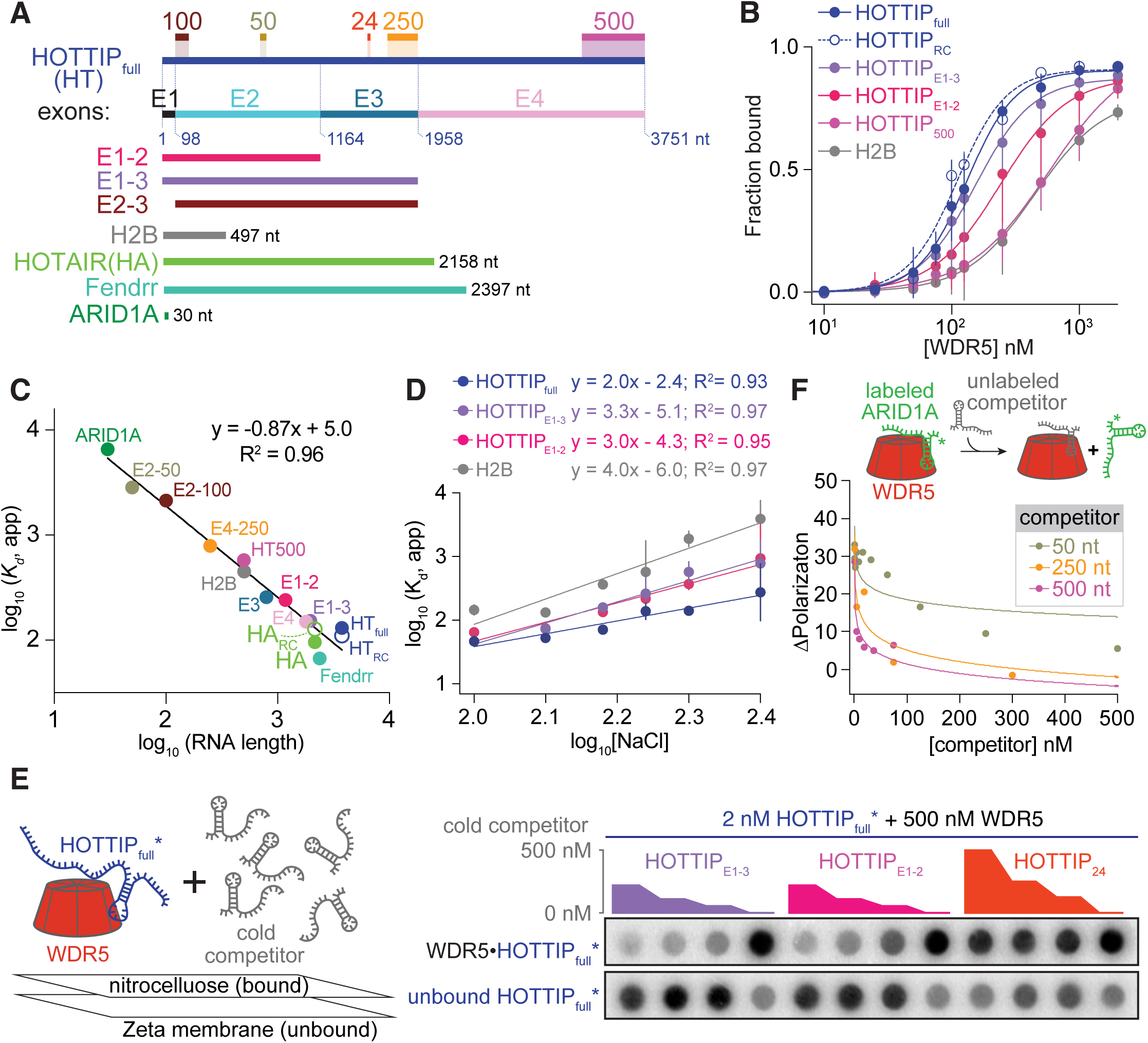
WDR5 binds RNA promiscuously *in vitro*. **(A)** Diagram of different RNA species used in the study including HOTTIP, HOTAIR, Fendrr, H2B, ARID1A, as well as fragments and reverse-complement sequences indicated along with their length in nucleotides (nt). **(B)** Equilibrium filter binding (FB) measurements showing complete binding curves for several different length RNA to WDR5. Shown are the average from at least three replicates ± S.D., further binding curves for other RNA in 2A are presented in Figure S2C. **(C)** Plot of average *K*_d,app_ versus the RNA length plotted with log-log scaling. Due to technical limitations of weak binding, fluorescence polarization was used to measure ARID1A (30 nt) and HOTTIP_24_ affinities. Nevertheless, these data lie on the linear regression fit of the FB data only, thus we conclude that the measurement modality has little influence on our reported affinities. (R^2^ and equation for the linear regression presented). **(D)** Plot of log (*K*_d,app_) against log [NaCl] for HOTTIP full length, HOTTIP_E1-3_, HOTTIP_E1-2_, and H2B RNA fit by linear regression (R^2^ and equation for the linear regression presented). **(E)** Depiction of competitive equilibrium filter binding assay (*left*) and measurements (*right*; nitrocellulose and Zeta-probe membranes, top and bottom panel, respectively) showing that HOTTIP_E1-3_ and HOTTIP_E1-2_ fragments are both able to competitively deplete radiolabeled HOTTIP_full_ RNA bound to WDR5. HOTTIP_24_ a 24 nt, non-binding RNA displays no competion at a concentration as high as 500 nM. The concentration of WDR5 is ∼4-fold > *K*_d,app_ for HOTTIP-WDR5 interaction. **(F)** Competitive fluorescence polarization measurements where unlabeled HOTTIP RNA fragments (HOTTIP_50_, HOTTIP_250_, HOTTIP_500_) were individually titrated against WDR5 (4 µM) with labeled ARID1A RNA (10 nM). See also Figure S2

Consistent with previous immunoprecipitation studies^42,44^ and strong enrichment in our WDR5-crosslinked RNA pool, we found that WDR5 indeed binds HOTTIP RNA directly (Figure 2B) with remarkably tight and modestly cooperative binding affinity (*K*_d,app_ 131 ± 9 nM; Hill coefficient of 2.3 ± 0.3). Native^75^ and various RNA refolding conditions did not appreciably alter the WDR5 affinity measured nor the apparent mobility in a native gel. Next, we tested the specificity of WDR5-RNA interaction. As prior affinity measurements of WDR5 binding of HOTTIP were qualitative, non-equilibrium measurements with the relatively short H2B or U2 RNA as the only negative controls^42,44^, legitimate questions remain as specificity of WDR5 for a specific RNA sequence. We started by screening varying lengths of *in vitro* transcribed HOTTIP RNA fragments, chosen by exon boundaries, yet agnostic of potential RNA sequence or domain structure (Figure 2A, Table S1). In addition to HOTTIP fragments, we examined four other distinct RNA species: 1.) HOTAIR RNA and fragments thereof – this RNA is thought to bind PRC2 complex, albeit with considerable promiscuity^51,58,76^, 2.) HOTTIP_RC_ and HOTAIR_RC_ – these are artificial RNA constructs that are the reverse complement of HOTTIP and HOTAIR, respectively; 3.) Fendrr RNA – this lateral-mesoderm specific lncRNA co-immunoprecipitated with WDR5 at levels similar to HOTTIP RNA^43^, and 4.) H2B RNA, employed as a negative control in previous pull-down assays^42,44^. We found that HOTAIR, HOTTIP_RC_ and Fendrr RNAs display similar WDR5-binding affinity (*K*_d,app_= 100 ± 10 nM, 110 ± 20 nM, and 67 ± 6 nM respectively) to that of HOTTIP RNA (Figure 2B, S2C). We note that unlike prior qualitative experiments (for which no experimental data was presented) ^42^, HOTTIP_E1-2_ does not account for the WDR5’s affinity for the full HOTTIP. Compared to the HOTTIP_full_, the H2B RNA has an ∼3.5–fold lower binding affinity (*K*_d,app_= 450 ± 50 nM), similar to an approximately length-matched fragment of HOTTIP (HOTTIP_500_) with *K*_d_,_app_ of 580 ± 60 nM (Figure 2B). The failure of any smaller fragment of HOTTIP to account for the WDR5’s affinity for the full HOTTIP and similar affinity of WDR5 to HOTTIP _RC_ and HOTTIP suggest the lack of a defined sequence motif or structural element for WDR5 binding. To explore the RNA length-dependence on WDR5 affinity, we expanded our equilibrium binding measurements to HOTTIP fragments of different lengths as well as other control RNAs described above – collectively probing 14 different lengths of RNA ranging from 24 to 3,764 nucleotides (Figure 2C, S2C). WDR5 binding was negligible for a 24nt fragment of HOTTIP within the concentration range examined by equilibrium filter binding assay (up to 2 μM) and fluorescence polarization assays (up to 60 µM), precluding accurate determination of dissociation constant (Figure S2C-D). For all larger RNA examined, we found that the binding affinity was almost perfectly-correlated to the length of RNA, with longer RNA displaying mid-nanomolar *K*_d,app_ values and shorter RNA ranging from high nanomolar to micromolar *K*_d, app_ values. A plot of RNA length versus *K*_d,app_ on a log_10_ scale shows a linear dependence with a slope of ∼ -1 (Figure 2C), despite being composed of a collection of different sequences. Using the 30-mer span of the smallest RNA-fragment with detectable binding as a minimal threshold for WDR5 binding, we performed a Monte Carlo simulation of how WDR5 binding affinity would scale for the different RNA-lengths in the absence of structure or sequence specificity and found an excellent recapitulation of the experimentally observed binding trend (Figure S2E). This linear correlation between length of RNA indicates promiscuity in RNA-binding by WDR5, reminiscent of the initially described RNA-binding by PRC2^58^. However, unlike the recent reports of higher affinity of PRC2 to RNA G-quads^59–61^, we observed no apparent WDR5 binding differences to G-quadruplex forming sequences as a function of stabilizing or refractory salts (data not shown).

As the RNA binding by WDR5 is only modestly sensitive to sequence and the electrostatic potential of WDR5’s surface is markedly electropositive^77–79^, we reasoned that coulombic interactions between the RNA-phosphate backbone and WDR5 surface patches composed of basic amino acid side chains could be a substantial component of the affinity. To examine the role of electrostatics in WDR5 binding to RNA, we systematically varied the concentration of monovalent salt in the binding conditions. The RNA-binding interface of WDR5 remained dependent on the length of the RNA (Figure 2D). As the slope of this plot is thought to be representative of the number of salt bridge interactions^58,80^, we infer that there are ∼2-4 salt bridges for WDR5-RNA interaction.

To examine whether the RNA-binding interface(s) on WDR5 are shared or overlapping between different RNA, we carried out competitive binding assays. We incubated [^32^P] body-labeled RNA with excess WDR5 (∼4-fold higher concentration than the *K*_d,app_), and titrated competing unlabeled “cold” RNA (Figure 2E). We found that both HOTTIP_E1-3_ and HOTTIP_E1-2_ at higher concentrations were successfully able to compete full-length labeled HOTTIP, indicating that all these RNAs share at least partially overlapping binding interfaces on WDR5. Reciprocal cold competition experiments with labeled medium-sized fragments revealed similar results (Figure S2F). As a negative control, a 24nt RNA (HOTTIP_24_) that does not show any appreciable binding to WDR5 (Figure S2C-D) was unable to compete with HOTTIP, HOTTIP_E1-3_ or HOTTIP_E1-2_ at concentrations as high as 500 nM (∼250-fold above labeled RNA concentration; Figure 2E and S2F). Using an orthogonal approach, we employed fluorescence polarization to probe competition between smaller-sized RNA fragments. WDR5 weakly binds to a 30nt RNA with micromolar affinity (*K*_d, app_ 7 µM, Figure S2C). The 30nt RNA was competed out by longer HOTTIP fragments (50, 250, 500 nt) (Figure 2F) in a length-dependent manner, as per competitive binding simulations (Figure S2G). Collectively, these experiments argue for at least partially charge-based binding of HOTTIP RNA fragments to a common WDR5 interface. The affinity of this WDR5 interface scales as a function of the RNA-length, rather than recognition of a particular sequence or structural element, with *in vitro* promiscuity observed congruent with cell-based measurements.

### Molecular dissection of the RNA-binding interfaces of WDR5

To map the RNA binding interface(s) of WDR5, we conducted mutagenesis studies based on the available structural data (Figure 3A) ^26,77–79,81^. Using non-equilibrium GST pull-down assays followed by RT-qPCR, previous studies identified F266A as a putative RNA-binding deficient mutant that was then deployed in mESCs to argue for the functional importance of RNA-binding for pluripotency^44^. Concerned that this F226A mutation targets the hydrophobic core of the protein rather than a surface residue that might plausibly make direct RNA contacts (Figure 3A), we sought to more closely examine its impact on RNA-binding using quantitative equilibrium affinity measurements. In contrast to the previous measurements, we found that the F266A mutant bound HOTTIP RNA with slightly higher affinity than WT WDR5 at 4°C, while at 37°C the F266A mutant displayed substantially eroded apparent affinity relative to WT (Figure 3B). This led us to suspect the stability of fold of the F266A mutant may be compromised at higher, physiologically relevant temperatures.

**Figure 3.**
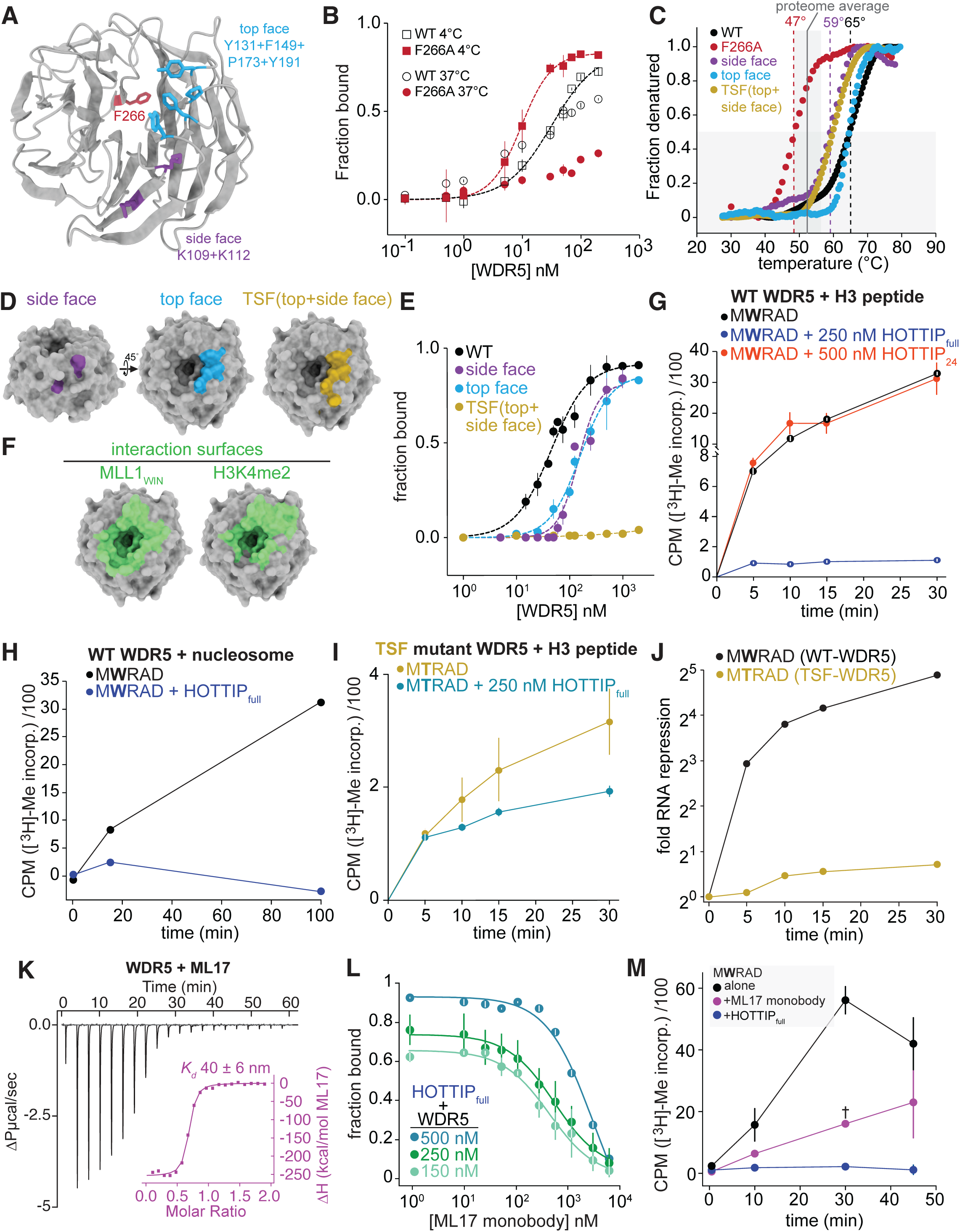
WDR5 has more than one RNA-binding interface. **(A)** Structure of WDR5 (PDB ID: 4ESG) indicating the mutated residues. **(B)** Equilibrium filter binding measurements of wild type (WT) and F266A mutant WDR5 to HOTTIP RNA at 4°C and 37°C. (*K*_d,app_ WT_4°C_ 30 ± 10 nM, F266A_4°C_ 10 ± 1 nM, WT_37°C_ 20 ± 20 nM, F266A_37°C_ > 200nM). Note that these preparations were performed side-by-side, and the WT protein furnished higher affinity than many of our other purifications. **(C)** The optical melting profile of WDR5, F266A, side face, top face and TSF (top + side face) mutants using differential scanning fluorimetry shows that TSF mutant is thermodynamically stable at the temperature of the equilibrium filter binding assays. The measured melting temperature (dotted lines) represents the point at which 50% of the protein is in an unfolded state. Estimated average and standard deviation of melting temperature of human proteome^86^ is also shown for reference. **(D)** Structure of WDR5 (PDB ID: 4ESG) indicating surface location of the mutated residues in the ‘side face’ mutant (K109, K112) and the ‘top face’ mutant (Y131, F149, P173, Y191). **(E)** Equilibrium filter binding measurements comparing the binding of the indicated variant of WDR5 to HOTTIP RNA. The TSF mutant incorporates mutations in the top face and side face mutants; see panel A. **(F)** Structure of WDR5 indicating the interacting surface of MLL1 WIN motif peptide (PDB ID: 4ESG) and H3K4me2 (PDB ID: 2H6N). WDR5 residues within 4.5 Å of any atom of the indicated binding partner are highlighted in green. Note that the CryoEM structure of the MLL1 KMTase core on H2Bub nucleosomes displays more extensive contacts (Figure S3F). **(G)** Time course measurements from histone methyl transferase assays measuring the transfer of the [^3^H]-labeled methyl group from S-adenosylmethionine to H3_1-20_ peptide by the MLL1 core HMTase complex (MWRAD: MLL1+WDR5+RbBP5+ASH2L+DPY30) in the absence or presence of indicated RNA. Independent triplicate time points indicated were quenched by spotting on phosphocelluose, then washed extensively and subjected liquid scintillation counting (counts per minute). Error bars represent S.D. **(H)** Same as (G) except that reconstituted nucleosomes^127^ were used as the substrate instead of the H3 peptide. **(I)** Same as (G) except that the TSF mutant of WDR5 was used instead of wild type in the MLL1 complex. **(J)** Comparison of the RNA-mediated repression of histone methyl transferase activity of MLL1 complex consisting of either WT or the TSF mutant WDR5. Fold repression calculated from data in (G) and (I). **(K)** Isothermal calorimetry thermogram and binding curve of ML17 titrated into WDR5 solution. **(L)** Equilibrium filter binding competitive measurements with 500, 250, and 150 nM WDR5 in the presence of HOTTIP RNA (<< 1 nM) with increasing concentration of the monobody. The IC_50_ is calculated to be 400 ± 70 nM, 500 ± 70 nM and 2600 ± 600 nM, at 150, 250, and 500 nM WDR5, respectively. Shown are the average from three replicates ± S.D. **(M)** Time course measurements from histone methyl transferase assays of MLL1 complex in the absence or presence of ML17 monobody (5 µM) or HOTTIP_full_ (250 nM) RNA under conditions as in panel 3G. Shown are the average from two replicates ± S.D. except for † (single replicate). See also Figure S3

We investigated the thermal stability of F266A using circular dichroism (Figure S3A), and found that F266A mutant has a much lower melting temperature (47°C with the earliest indicia of unfolding detectable at 40°C) than the WT protein (65°C). Similar results were obtained using differential scanning fluorimetry (Figure 3C). Our data suggest that the reported apparent RNA-binding deficiency of F266A mutant, and likely its physiological consequences in mESCs^44^ may be independent of RNA-binding capacity, but rather depend on its thermodynamic meta-stability, that yields unfolded protein as a function of time at physiological temperature. This WDR5 destabilization is also supported by findings from a recent study that showed that the F266A mutation leads to a reduced apparent affinity with a peptide that binds to WDR5^82^, far from the site of this mutation. Consequently, further investigation using mutants with sufficiently high thermal stability to avoid this pitfall was required to map the RNA-binding interface of WDR5. To this end, we examined a panel of WDR5 surface-residue mutants (Figure S3B). Initially we focused on making neutralizing mutations to a series of positively charged basic patches of WDR5, due to the observed electrostatic contribution (Figure 2D) and dominant roles played by Arg/Lys in ribonuclear-protein structure meta-analyses^83–85^. We found one region in particular, that resulted in a 3-5-fold RNA-binding deficiency (Figure 3D-E and S3C-D). This mutant (‘side face’: K109A+K112M) has a melting point of 59°C slightly below the WT protein (Figure 3C and S3A), suggesting that the modest erosion of binding is due to interface perturbation rather than thermally instability. Although this mutant diminished apparent RNA-binding, the decrement was limited, even when deployed in combination with mutations of three other adjacent residues on the radial surface of WDR5 (Figure S3C-D).

Relative insensitivity to incremental point mutations on the radial surface of the protein compelled us to examine the top face of WDR5, known to be the site of mutually exclusive engagement of the WIN (*W*DR5 *In*teracting) motif of the MLL1 polypeptide and histone H3 (Figure 3F) ^25,77–79^. We found that a tetramtuant of this surface (‘top face’: Y131A, F149A, P173A, and Y191A, Figure 3A and 3D) also showed a modest ∼3-fold erosion in binding affinity (Figure 3E) indicating that at least two discrete surfaces on WDR5 are important for RNA binding or a large continuous patch spans both sites. Consistent with this hypothesis, combining both sets of surface mutations (“TSF”, Figure 3D) completely abrogated detectable WDR5-RNA binding in the concentration regime examined (Figure 3E). Despite 6 amino acid changes, the TSF WDR5 mutant remains reasonably thermodynamically stable with a melting temperature of 58 °C (Figure 3C, S3A), well above the average human protein melting temperature estimates^86^. Although further points of contact cannot be ruled out, our targeted mutation analyses indicate that energetically critical contacts to RNA include both the WDR5 top face and side face of the second propeller blade. Interestingly, structural prediction using AlphaFold3^87^ also predicts similar surface of WDR5 to engage with RNA (Figure S3E), supporting our findings that RNA interacting surface of WDR5 overlaps with that of WIN motif.

### RNA-binding to WDR5 markedly inhibits the histone methyltransferase activity of the MLL1 complex

As our mutagenesis data indicate that the RNA-binding interface on WDR5 has some overlap with the MLL1-binding region (Figure 3F, S3F) known to be important for the catalytic activity^25,88^, we examined whether RNA-binding by WDR5 can impact the MLL1 complex’s catalytic activity. We conducted KMTase assays in the presence of RNA using purified components of the core MLL1 KMTase complex^24,89^. Surprisingly, we found that the KMTase activity of MLL1 was diminished by >15-fold upon the addition of full length HOTTIP RNA at near-saturating concentrations (Figure 3G). This inhibition of the MLL1 complex’s enzymatic activity is RNA concentration dependent around the complex *K*_d_ (Figure S3G). The same impact on activity was not observed with a 24nt RNA (HOTTIP_24,_ Figure 3G), which displays no detectable binding to WDR5 at the 500 nM concentration used (Figure S2C-D), consistent with the hypothesis that RNA binding by WDR5 is responsible for this diminution in activity. In addition to the H3 peptide substrate, the RNA-mediated inhibition of MLL1 complex KMTase activity was also observed using the nucleosomal substrates (Figure 3H). To confirm the interpretation that the decrease in MLL1 activity was a consequence of RNA-binding to WDR5, and not a non-specific effect of RNA addition or potential RNA-binding by another subunit, we conducted KMTase assays with an MLL1 core KMTase complex with several perturbations to the WDR5 subunit. In particular, we performed side-by-side histone methyltransferase assays with an MLL1 core KMTase complex with WT WDR5 (MWRAD), a complex lacking WDR5 (MRAD), and a complex where WDR5 was supplanted with the TSF mutant that completely abolishes RNA-biding (MTRAD, Figure 3E). As previously observed, absence of WDR5 severely diminishes MLL1 KMTase activity^24,26,89,90^, although there is little discernable impact on the activity for this subcomplex with HOTTIP_full_ at concentrations that inhibit the complex with WDR5 (Figure S3H). The TSF WDR5 mutant complex (MTRAD), despite its reduced activity as compared to MWRAD, is still more enzymatically active than the complex lacking WDR5 (Figure 3I). Upon addition of equimolar HOTTIP_full_ to MTRAD, there is a slight decrease in KMT activity (Figure 3I), but this decrement is far more modest than the effect of HOTTIP_full_ addition to MWRAD (Figure 3J). Furthermore, titration of HOTTIP_full_ seems to have no effect on the KMT activity of MTRAD (Figure S3H), supporting the interpretation that the RNA’s ability to bind WDR5 is the crucial determinant of the KMT activity repression.

To further interrogate the RNA-binding interface on WDR5 using an orthogonal non-invasive approach, we sought to design proteins that might interfere with the RNA-binding capacity of WDR5. We generated short (∼12 kDa) synthetic proteins on an FN3 scaffold, called monobodies^91^ with mid-nanomolar binding affinity for WDR5. First, we tested our previously described “S4” monobody that binds to the top face of WDR5^92^ (Figure S3I). Although S4 monobody disrupts RNA binding (Figure S3I), it also binds to RNA directly precluding interpretation of competitive binding to a shared WDR5 interface. To overcome this limitation, we engineered ML17 another monobody that binds to WDR5 with mid-nanomolar affinity (Figure 3K) but does not bind RNA even when incubated in concentrations as high as 10 μM – beyond the upper limit we deploy in WDR5-RNA binding assays (Figure S3J). Upon titrating ML17, in the presence of HOTTIP and WDR5 under conditions where we normally detect robust binding, we found that ML17 is able to compete with RNA (Figure 3L). This effect is not exclusive to full-length HOTTIP as we found that ML17 is also a successful competitor of WDR5 complexes with fragments of HOTTIP (Figure S3K), consistent with the RNA binding promiscuity of WDR5 being due, at least in part, to a common RNA-binding surface. Thus, ML17 can be used as an additional modality to interrogate key features of the RNA binding surface of WDR5.

Next, we tested whether the portion of the WDR5 RNA-binding surface that overlaps with that of ML17 binding is critical for MLL1 KMTase activity. The addition of saturating amounts of ML17 monobody modestly reduced MLL1 KMTase activity, indicating that the ML17-senstive portion of the RNA binding interface to WDR5 is also important for MLL1 enzymatic activity (Figure 3M). These data also argue that there is no specific requirement for RNA in this mode of KMTase inhibition, as a protein can achieve similar inhibition, presumably by competition for the similar portions of WDR5-surface needed for MLL1 complex activity^25,88^. However, the effect of ML17 KMTase inhibition is far less dramatic than that of HOTTIP RNA, despite a greater degree of saturation in binding (2 orders of magnitude versus 2-fold above *K_d_*), suggesting that it represents a partial effect. This could be due to a smaller footprint of ML17 than HOTTIP RNA or sampling of non-identical surfaces which contribute to KMTase activity to different extents. Collectively, these findings are consistent with the model that binding of sufficiently long RNA by WDR5 competes for catalytically important interfaces with the MLL1 complex WIN motif binding site, culminating in impaired methyltransferase activity. Yet all the evidence for this model of interface competition is indirect.

### WDR5-RNA binding disrupts MLL1 core complex stability

We sought to understand the molecular mechanism by which WDR5 binding of RNA impairs the histone methyltransferase activity of the complex and explicitly test the model that RNA competes for the critical MLL1-WDR5 interface. As several of the WDR5 interfaces to core MLL1 complex subunits have been demonstrated to be mutually exclusive with structurally defined sites of interaction with other engaging factors^93–98^, the complex may be somewhat more dynamic under physiologic conditions^98,99^ than the typical monolithic view^18,24^. The WDR5 mutations that we found to cause the most severe disruption in RNA-binding were along the top face, where MLL1 and histone tails bind to WDR5, and a positively charged side face of the second propeller blade (Figure 3D, 3F). Given that regions of top surface of WDR5 are responsible for mutually exclusive binding of MLL1 that is required for HMTase core complex stability and activity^26,89,93,100^, we reasoned that RNA-binding by WDR5 could impair HMTase activity of MLL1 complex competitively by disrupting the MLL1-WDR5 protein-protein interface. We performed an *in vitro* pull-down assay to test the impact of RNA on the binding of WDR5 to GST-tagged MLL1 C-terminal fragment immobilized on glutathione beads (Figure 4A). Addition of HOTTIP_full_ RNA led to a concentration-dependent reduction of WDR5-MLL1 interaction (Figure 4A, 4B, S4A, S4B). Free NTPs at a concentration that was per nucleotide molar- and composition-matched to HOTTIP_full_ (3754-fold the indicated concentration, Figure 4A, 4B, S4A, S4B) or the non-binding HOTTIP_24_ (Figure S4C) had no effect on WDR5-MLL1 interaction, consistent with competition originating from WDR5-binding RNA and ruling out any non-specific effect of an RNA-mediated increase in the ionic strength of the binding conditions.

**Figure 4.**
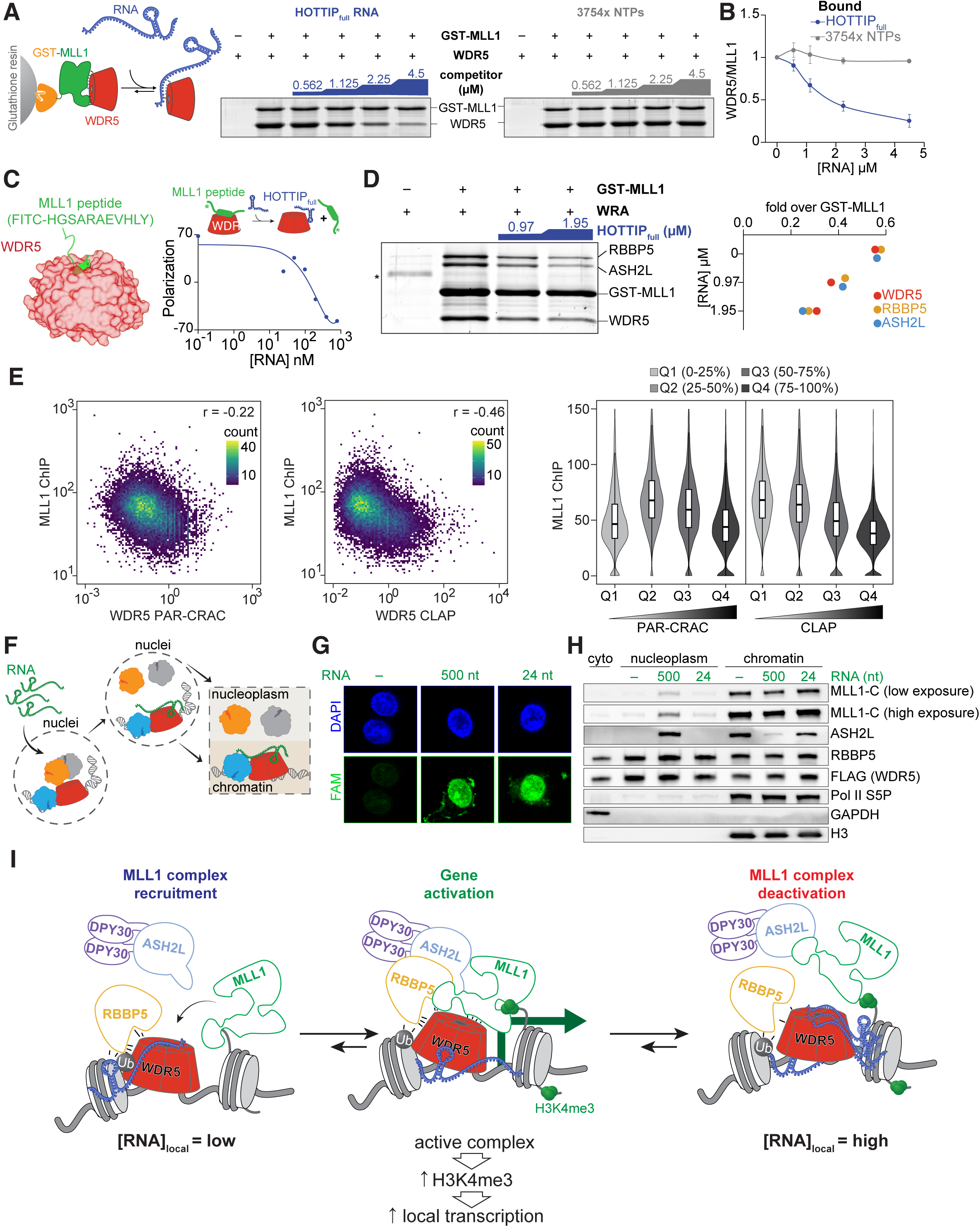
RNA competitively displaces MLL1 polypeptide from WDR5. **(A)** *Left*, Schematic of GST-Pull down assay to test the role of RNA in the binding of WDR5 to GST-MLL1 immobilized on glutathione beads. *Right*, SYPRO-Ruby staining of GST-MLL1 and WDR5 proteins eluted from glutathione resin after GST-MLL1 and WDR5 incubation in the presence of the indicated concentration of HOTTIP full length RNA (HOTTIP_full_, *left*) or free NTPs, concentration-matched per-nucleotide to HOTTIP (3754x the indicated concentration, *right*). **(B)** Quantitation of WDR5 bound to GST-MLL1 across the indicated range of RNA/NTP titrations. Shown are the average from two replicates ± S.D. **(C)** *Left*, Structure of WDR5 bound to the MLL1 peptide (PDB ID: 4ESG). *Right*, Competitive fluorescence polarization assay of labeled MLL1 peptide (10 nM) and WDR5 (250 nM) with increasing concentrations of HOTTIP full-length RNA. **(D)** *Left*, SYPRO-Ruby staining of the MLL1 complex subunits bound to immobilized GST-MLL1 under the indicated concentration of HOTTIP full-length RNA. * Uncharacterized protein fragment from RBBP5 protein prep. *Right*, Quantitation of gel staining for the indicated proteins. **(E)** Scatter plot of input normalized MLL1 occupancy (within 1kb upstream of the open reading frame) versus WDR5 PAR-CRAC counts (*left*) or WDR5 CLAP counts for transcripts that emanate from a given MLL1-bound promoter (*middle*) in HEK293 cells. r, Spearman correlation coefficient. *Right*, Distribution of input normalized MLL1 chromatin occupancy (within 1kb upstream of the open reading frame) as a function of WDR5-RNA interaction. Genes are ranked into quartiles based on WDR5 PAR-CRAC or WDR5 CLAP signal (Q1 to Q4: lowest to highest signal). Violin plots show the distribution of signal within each bin; embedded box plots show median (horizontal line) and interquartile range (box boundary). **(F)** Schematic for experiment presented in G and H, to test the RNA-induced disassembly of the MLL1-complex in cells. Freshly isolated nuclei from HEK293 cells were incubated with fluorochrome-labeled RNA (1 µM), followed by nuclear extraction to chromatin and nucleoplasm fraction. **(G)** Micrographs of representative nuclei showing the uptake of the indicated fluorochrome-labeled RNA. The 500nt RNA is a fragment of HOTTIP RNA that binds to WDR5 (*K_d_* 580 ± 60 nM); the 24nt RNA is a fragment of HOTTIP RNA that does not show detectable binding to WDR5 and is used as a negative control. **(H)** Representative western blot showing the partitioning of indicated proteins in the cytoplasm (cyto), nucleoplasm and chromatin fraction after fractionation with no RNA or a 500 nt fragment of HOTTIP RNA (1 µM) or a 24 nt fragment of RNA (1 µM). Note the increase of the MLL1-C and ASH2L subunits of the complex in the nucleoplasm fraction with corresponding decrease in the chromatin fraction by the 500 nt fragment that binds WDR5. RBBP5 and WDR5 equilibria do not appear to change in response to *in situ* RNA addition. No such change was observed for the 24 nt fragment RNA that does not bind to WDR5. MLL1-C denotes the C-terminal fragment of MLL1. Pol II S5P denotes serine 5 phosphorylated RNA Pol II. **(I)** Proposed model of RNA-mediated negative feedback regulation of MLL1 complex activity by local assembly/disassembly in response to local RNA concentration. Chromatin-bound RNA may promote recruitment of subunits/submodules of the MLL1 complex. With sufficiently higher local concentration of MLL1 (perhaps via other known chromatin interactions) relative to RNA, the complex assembles to a catalytically active form, displacing RNA from the WDR5 interface, leading to methylation of H3K4, which in turn, potentiates transcriptional activation. Highly abundant local RNA product of this transcription then competes with the MLL1-WDR5 interface, leading to the disassembly of the MLL1 complex, constituting a negative feedback loop to tune gene expression. See also Figure S4

While the competition between the C-terminal fragment of MLL1 and HOTTIP RNA indicates mutually exclusive binding to overlapping surfaces of WDR5, the MLL1 protein interface is substantial (Figure S3F). To further refine the region within this MLL1 interface that impacts RNA binding, we probed the short MLL1 peptide that represents the minimal *W*DR5-*<u>in</u>*teracting (“WIN”) motif (Figure 3F, 4C^96,100^) via competitive fluorescence polarization of labeled MLL1 WIN peptide in the presence of HOTTIP_full_. Indeed, this MLL1 WIN peptide can be competitively displaced from WDR5 by full length HOTTIP (Figure 4C), indicating overlapping binding surfaces, consistent with residues that impact RNA binding include those that directly interact with this peptide (Figure 3E). We note a similar *K_d_* for MLL WIN peptide using fluorescence polarization (Figure S4D) to that previously reported^96,100^, affirming the validity of this assay.

To further refine the portion of the MLL1-WDR5 interface that can be competitively antagonized by RNA binding, we sought to query the central cavity of WDR5 (Figure S4F) that harbors the arginine binding pocket^95,96^. We examined whether the small molecule WDR5 inhibitor MM-401 that engages only the central portion of the MLL1 WIN/H3 peptide binding surface, particularly the arginine-binding declivity^88^, could compete with RNA binding. We find titration of MM-401 causes no apparent decrease in RNA-binding up to 10 μM (Figure S4E), well above the subnanomolar *K_d_* for the molecule^88^. MM-401 poses far less steric demand to the top surface of WDR5 than the WIN motif peptide or larger MLL1 protein interface (Figure S4F). Thus, we conclude that unlike the MLL1 protein contacts to WDR5, the WDR5 central pocket is not an energetically important contributor to the RNA interface, whereas the coronal top surface that surrounds this cavity, is important for both MLL1 and RNA engagement. Next, we tested the impact of RNA on the assembly of the MLL1 core KMTase complex. Consistent with the critical role of the WDR5-MLL1 interaction in the assembly of MLL1 complex^24–26,90,99^, addition of HOTTIP_full_ led to a reduction in the stable binding of WDR5, RBBP5 and ASH2L subunits with immobilized MLL1 (Figure 4D). These data indicate that RNA binding to WDR5 can lead to disassembly of the MLL1 complex.

Next, we sought to test RNA-mediated disassembly of MLL1 complex in cells. Our model predicts a reduction in the chromatin occupancy of MLL1 with the increase in WDR5-RNA interaction at a given promoter. Despite the limitations of the redundancy between MLL1 complexes, steady state levels of gene expression across a large cell population, and presumed asynchrony of the putative MLL1 complex assembly/disassembly cycle, we observe a negative correlation between MLL1 chromatin occupancy and WDR5-RNA interaction in cells (Figure 4E), consistent with our model. To empirically test RNA-mediated disassembly of the MLL1 complex in cells, we incubated nuclei with fluorochrome-labeled HOTTIP_500_ and HOTTIP_24_ RNA followed by subnuclear fractionation into nucleoplasm and chromatin pools (Figure 4F, 4G). Strikingly, we observed an increase in the nucleoplasm pool of the MLL1-C and ASH2L subunits of the complex with a corresponding decrease in the chromatin fraction when nuclei are incubated with the HOTTIP_500_ RNA that binds WDR5 (Figure 4H, S4G). No such change was observed for the HOTTIP_24_ RNA that does not bind to WDR5. Notably, WDR5 and RBBP5 distributions remain unchanged under these conditions, consistent with chromatin tethering of these subunits by *cis*-acting local RNA.

Taken together, our data indicate that RNA binding to WDR5 competes with its protein-protein interface with MLL1 leading to potential disassembly of the MLL1 complex. As this interface is particularly important for MLL1 activity^88,96^, this finding provides a mechanistic basis for the RNA-mediated inhibition of MLL1 complex KMTase activity.

## DISCUSSION

RNA is posited to be a central player in chromatin regulation, actively engaging with transcription factors and remodeling complexes^3–5,7,8^, yet the mode of interaction, specificity determinants, and functional outcome remains contentious. In this work, we systematically defined the molecular details of direct WDR5 binding of RNA and its dramatic consequences on histone methyl transferase activity of the MLL1 complex. Our *in cellulo* and *in vitro* investigations show that WDR5 binds tightly and directly to a multitude of RNAs without any apparent sequence specificity. *In vitro*, this promiscuity manifests as WDR5 engagement with RNA in a length-dependent manner and we have mapped critical determinants of this binding. We show that RNA competitively inhibits the interaction of WDR5 to MLL1 polypeptide, leading to disassembly or a conformational change that inactivates the histone methyltransferase activity of the MLL1 complex.

### Promiscuity in WDR5-RNA interaction

A burgeoning body of literature has proposed that RNA, via notionally specific direct binding to the WDR5 subunit, recruits the MLL1 complex to specific loci in the genome where it then acts to install the H3K4me3 mark^42–44,46–48^. Yet prior to this work, the specific WDR5-RNA recognition that this model requires has not been unambiguously examined *in cellulo* nor been subjected to rigorous quantitative equilibrium binding studies. Further uncertainty has been cast on these models by recent claims that apparent RNA binding to WDR5 and several other transcription-associated factors, is an artifact stemming from low stringency of the corresponding cell-based cross-linking and complex isolation methods as well as normalization choices^62^. Using our denaturing PAR-CRAC technique that encompasses stringent wash conditions to minimize non-specific binding, we identified a large cohort of RNAs that directly crosslink to WDR5 in proportion to their abundance, agnostic of the underlying sequence. We note that data from our PAR-CRAC analysis and recent CLAP analysis both indicate no sequence specificity in WDR5-RNA binding. The divergence in interpretation from the two datasets and analyses may originate from thresholds to call a bona fide binding. In the absence of a defined sequence preference that culminates in an observable sequence motif, making this distinction from these types of data alone is particularly challenging.

To overcome these challenges and further investigate the nature of WDR5-RNA interactions, we conducted a series of complementary *in cellulo* and *in vitro* assays that further attest to the biological importance of WDR5 binding RNA. First, our *in situ* fractionation coupled to RNase treatment revealed that the chromatin occupancy of a considerable fraction of WDR5 is sensitive to RNase. We note that this solubilization-based method to test RNA-dependent occupancy of WDR5 is free of previously reported artifacts due to crosslinkers^61,65,101,102^ as well as RNase-triggered variations in chromatin immunoprecipitation^103,104^. Next, the lack of sequence agnostic yet direct binding to RNA is well-supported by our *in vitro* biochemical characterization. The quantitative binding experiments show direct binding of HOTTIP RNA to WDR5 with mid nanomolar affinity which is well within the cellular concentrations of WDR5 (∼170 nM^105^ compared to a *K_d_* of 130 nM) and RNA. Moreover, the affinity of WDR5 to RNA is similar to that of protein-protein interactions of the MLL (Figure S2H) complex subunits, other chromatin modifying complexes^106^, and sequence-specific RNA-binding proteins^107,108^, arguing for the functional relevance of the WDR5-RNA interaction. Although HOTTIP RNA was especially enriched in our PAR-CRAC studies, it afforded comparable affinity to other similarly sized RNA species when tested *in vitro*. Rather than any apparent sequence specificity, we found that the affinity of the WDR5-RNA engagement *in vitro* appears to be a perfectly linear function of the length of RNA. Despite sampling several different RNA folding conditions for HOTTIP including “native” preparations^75^, we cannot rule out the possibility that the HOTTIP we deployed in these equilibrium binding studies may not recapitulate the element of the *in vivo* structure critical for specific recognition. While HOTTIP represents an outlier in our PAR-CRAC data, the vast majority of these data also support promiscuous RNA binding to the complex nuclear pool of RNA, manifesting more as a function of nuclear abundance. The cell-based PAR-CRAC data and *in vitro* binding assays reflect WDR5’s avidity for a small nonspecific segment of RNA, presumably an ∼ 30nt span (the smallest RNA with detectable binding to WDR5). That avidity can scale with potential binding sites iterated along the length of a single RNA when it is present in isolation (as in the *in vitro* binding assays) or as a function of site abundance within a crowded RNA population in the nuclear milieu (as in our PAR-CRAC data). The length-dependent binding we observe *in vitro* also implies that the previously reported reduced apparent affinity of WDR5 to H2B RNA versus HOTTIP in a qualitative format^42,44^ is indeed due to the order of magnitude shorter length of this RNA rather than sequence specificity that was proposed. Additionally, our competitive binding assays show that RNAs bind to an overlapping interface on WDR5 without sequence specificity. The lack of sequence specificity in WDR5-RNA interaction bears resemblance to previous reports on promiscuous PRC2-RNA interaction^56–58^. However, recent studies argue that PRC2 has higher affinity for G-rich RNA sequences, especially those that form RNA G-quads^59–61^. We did not find such preference for RNA G-quads for WDR5-RNA interaction, possibly reflecting on the differences in the genomic contexts in which these two HMTase complexes function. The accumulated weight of our *in cellulo* and *in vitro* evidence supports the conclusion that *bona fide* and functionally relevant WDR5-RNA interactions occur in cells without any strict sequence or structure specificity via a common RNA binding interface.

### The RNA-binding interface of WDR5

Although WDR5 is widely considered to be an RNA binding protein^42–48^, the surface with which it binds to RNA has remained elusive. Previous studies showed that the F266A mutant of WDR5 is deficient in RNA-binding by qualitative measures at an indeterminate temperature^44^. However, our quantitative equilibrium binding measurements displayed no RNA-binding deficiency compared to WT protein at 4°C. Only at 37°C did we detect lower apparent affinity for this mutant, attributable to its lower thermostability as opposed to an RNA binding affinity decrement of the folded protein at physiological temperature. The low thermal stability of F266A can perhaps be attributed to the deeply buried location of the F266 residue in the hydrophobic core of WDR5 where it stabilizes inter-blade interactions within the WD40 propeller structure. To obtain a thermodynamically stable mutant with RNA-binding deficiency, we designed multiple mutants focusing on the positively charged patches of the electrostatic surface and the MLL/histone binding interface. We found that a mutant targeting several of these surfaces (TSF mutant) had the most deleterious effect on RNA-binding. The engagement of the top face of WDR5 with RNA is consistent with top face of WD40 proteins being a hotspot of WDR5s interactions^88,93,96,109^. Whereas the involvement of lysine residues is consistent with the substantial coulombic component of many protein-RNA interactions^85^ and our data indicating several interfacial salt bridges.

Our ensemble of point mutants and competitive binding assays presents a fine-grained map of the RNA-binding interface of WDR5. First, the WDR5 residues in the ‘top face’ mutants play pivotal role in interaction with the WIN motif of MLL1 – F149, Y191, and P173 form a hydrophobic pocket to interact with H3769 of MLL1^93,96^ while Y131 may provide a complementary hydrophobic cage to stabilize the extensive hydrophobic and cation–π interactions of F133 with the guanidino group of R3765 in the MLL1 Win motif^88,93^. The engagement of the top face is also confirmed by RNA-mediated competitive displacement of MLL1 WIN peptide from WDR5. However, the MM401 small molecule, which binds to the central pocket of WDR5 with a much smaller footprint than the MLL1 WIN peptide^88^ showed no competition with RNA. This suggests that, at the top face, RNA predominantly binds to the outer corona. Second, we observed that in addition to the corona of the top face, residues on the radial surface of WDR5 also contribute to RNA binding, and the contribution of both of these surfaces is clearly cooperative. These include K109 and K112 on the second propeller blade on a solvent-exposed surfaces, away from any known interacting partners. Distal to this site, another residue that modestly contributes to RNA binding is K250. Interestingly, K250 is located close to the WBM (WDR5-binding motif) site via which RBBP5^94^ and c-MYC^97^ are shown to interact with WDR5, suggesting that RNA may impinge upon WDR5-RBBP5/MYC interaction. Collectively, it is evident that WDR5 engages with RNA via multiple patches which could constitute a contiguous interface. Notably, this interface is shared across different RNAs as we observed length-dependent competitive displacement spanning different RNAs. We do not exclude the possibility that additional sites beyond those tested here may also contribute to RNA binding. Nevertheless, presence of multiple RNA-binding surfaces in disparate contexts with respect to MLL1 complex protein-protein interfaces may permit alternate modalities of RNA-mediated regulation.

### How the RNA-binding interface of WDR5 regulates MLL1 complex activity

Perhaps our most surprising finding is that RNA inhibits the histone methyltransferase activity of the MLL1 complex, as it appears to be in conflict with the previously ascribed role of RNA in promoting H3K4me3 installation^42,44,46,47,49,50^. The RNA-mediated inhibition of the MLL1 complex is WDR5-dependent, as the fold repression of activity upon addition of RNA was far lower in the complex lacking WDR5 or in the complex containing the RNA-binding deficient TSF mutant. Furthermore, RNA-mediated inhibition of the complex occurs above the WDR5 *K*_d_ for RNA, but is minimal below it. As an orthogonal approach, we developed the ML17 monobody as a tool to probe the RNA-binding interface of WDR5 to alleviate any concerns of thermal instability associated with mutagenesis. ML17 binds WDR5 with a 40 nM *K_d_*, does not bind RNA, and competes with RNA to bind to WDR5 (IC_50_ 250 nM) confirming that it can be used as at least a partial RNA mimic in terms of the WDR5 surface it occupies. ML17, like RNA, also inhibits the KMTase activity of the MLL1 complex, albeit far more modestly (5-fold at 30 minutes versus 30-fold for RNA) at super-saturating concentrations (125-fold above *K_d_*, versus 2-fold above for HOTTIP RNA). Collectively, these observations argue that the RNA-binding interface of WDR5 is mutually exclusive with an interface or conformation of the MLL1 core complex necessary for full KMTase activity.

A portion of the RNA binding interface of WDR5 that we describe here, especially the top face, is implicated in WDR5’s interactions with a plethora of proteins such as MLL1-4, SET1A-B^77,93^, H3 tail^79^, PDPK1^110^, a microprotein EMBOW^111^, and MBD3C^112^. Some of these interactions have been directly demonstrated to be mutually exclusive – MLL1 can compete with H3K4me2^79,113^. Similar competition can be inferred for others^77,93^ due to spatial overlap of engagement with the energetically critical arginine-binding pocket. Reminiscent of this mutual exclusivity, we report that RNA binding of WDR5 competitively inhibits the interaction between the SET domain-containing C-terminal region of MLL1 and WDR5. The full MLL1 polypeptide has been shown to make contacts with WDR5 at multiple surfaces^23^, but the most crucial is the top face where multiple residues in the WIN motif of MLL1 interact with WDR5^96^, and this interface is crucial for the structural integrity and activity of the MLL1 complex^26,89,93,114,115^. Consistent with this, we observed that RNA-mediated inhibition of WDR5-MLL1 interaction hinders the assembly of the MLL1 complex, providing a mechanistic basis of RNA-mediated repression of MLL1 activity. Based on our findings, we propose that RNA plays a key role in the maintenance of MLL1

KMT marks by contributing to the chromatin occupancy of WDR5 and concentration-dependent disassembly of the complex (Figure 4I). RNA-mediated chromatin tethering of WDR5 may increase its local concentration at regions of the genome that have basal transcriptional activity or otherwise bound RNA. While it is hard to imagine how the non-specific binding energetics that we find for WDR5 could impart RNA specificity to the recruitment process, they could increase the residence time of the WDR5 and RBBP5 subcomplex assembly on H2Bub nucleosomes, postulated to be an intermediate of MLL1 core complex assembly on chromatin^98^. Of note, the RBBP5-WDR5 interface is the weakest pairwise interface in the core MLL1 KMTase complex^89^, so its stabilization on chromatin could be plausibly rate limiting. Positive correlation of WDR5 PAR-CRAC (Figure 1C) and CLAP (Figure S1G) with WDR5 ChIP signal is consistent with RNA-mediated stabilization of this subunit at the site of its transcription and transcription-dependent recruitment of the H3K4me3 machinery as has been noted in several systems^41,116,117^. At low local RNA concentrations^118^, RNA could facilitate this process by affording additional binding energy, but be subject to competition with MLL1^C^ polypeptide recruited locally by multivalent binding to additional chromatin elements that can impart some locus specificity^21,36,39,119–121^, followed by ASH2L-(DPY30)_2_ assembly^98^. The high local concentration of MLL1 relative to RNA could favor the formation of catalytically competent complex to install H3K4me2/3 in promoter-proximal nucleosomes to promote transcription via several proposed mechanisms^14,122–126^. As local RNA concentration increases as a function of transcriptional bursting, disassembly of MLL1 from WDR5 becomes more favorable, leading to the feedback inhibition of the H3K4me3 activity. Different RNA-binding affinities, stemming from the length or amount of RNA, could determine the efficiency of displacement of MLL1 from complexes at target loci depending on RNA’s ability to compete for the WDR5 interface.

While other SET-domain proteins (MLL2-4, SET1A-B) interact with the top face of WDR5, with different affinities^96,100^, structural conformations^96,100^, and functional ramifications^114^, it remains to be tested whether and how RNA will impact WDR5s interaction with those proteins. As different members of the MLL family of complexes vary in terms of the dependence, affinity and structural aspects^100^ of WDR5’s interaction with the corresponding catalytic subunits, WDR5-RNA interactions may present another layer of differential regulation of the key KMTase complexes. Future studies will be focused on understanding the impact of WDR5-RNA interaction on the function of all these WDR5-containing complexes.

The identification of RNA as a modulator of the MLL1 complex challenges the canonical view of WDR5 as purely a scaffolding protein. Instead, WDR5 emerges as a multifunctional hub, balancing its interactions between RNA and protein partners. This regulatory mechanism could provide a layer of control over histone methylation, potentially integrating transcriptional activity with chromatin state. This mechanism may represent a broader paradigm in chromatin regulation, where local RNA abundance as a function of transcriptional activity provides a self-limiting signal for fine tuning the activity of chromatin-modifying complexes.

## RESOURCE AVAILABILITY

### Lead contact for reagent and resource sharing

Further information and requests for resources and reagents should be directed to and will be fulfilled by the lead Contact, Dr. Alexander Ruthenburg.

### Materials availability

Reagents generated in this study are available from the lead contact upon request.

## ACKNOWLEDGEMENTS

We thank the University of Chicago Functional Genomics Facility for Illumina sequencing. We thank Yali Dou for providing plasmids for SUMO-RBBP5 expression. We thank Michael Cosgrove for providing plasmids for GST-MLL1, His_6_-ASH2L and His_6_-DPY30 expression. We thank members of the Ruthenburg lab for helpful suggestions to improve the manuscript. This study was supported by the American Cancer Society (130230-RSG-16-248-01-DMC to A.J.R.), the Cancer Research Foundation (CRF) Fletcher Scholar, and NIH (R35-GM145373 to A.J.R, R01-DA036887 to S.K).

## AUTHOR CONTRIBUTIONS

Conceptualization: A.S.K., P.S., A.J.R; Methodology: A.S.K., P.S. and A.J.R.; Formal Analysis: A.S.K. and P.S. Investigation: A.S.K., P.S., M.S.W., A.G., A.K.; Writing – original draft: A.S.K.; Writing – review and editing: A.S.K. and A.J.R with input from all authors; Visualization: A.S.K. and A.J.R.; Supervision: S.K. and A.J.R; Funding Acquisition: A.J.R.

## DECALARATION OF INTERESTS

The authors declare no competing interests.

## SUPPLEMENTARY MATERIALS

Document S1: Figures S1 to S4

## STAR METHODS

### EXPERIMENTAL MODEL AND SUBJECT DETAILS

HEK293-FRT cell lines were grown in (DMEM + 10% FCS) at 37°C.

## METHOD DETAILS

### PAR-CRAC (*<u>P</u>*hoto-*<u>A</u>*ctivatable *<u>R</u>*ibonucleoside-enhanced *<u>C</u>R*ossIinking *a*nd analysis of *c*DNAs)

A FLAG-HA-6xHis-WDR5 pCDNA5 construct was incorporated into the FRT site of HEK293-FRT line (Invitrogen) using FLP-recombinase and Hygromycin B at 100 µg/ml for two weeks to select a monoclonal cell line. The HA and 6xHis tags permit tandem purification, including a His-Cobalt IP with a stringent denaturing buffer. The FLAG tag was used for direct western blot probing with anti-FLAG-HRP. 2 x 10 cm plates of this line were cultured to 75% confluency at 37°C, then 1 µl of 1 M 4-thiouridine (4sU, Sigma, re-suspended in DMSO) was added to the growth media (DMEM + 10% FCS) in each plate (100 µM final concentration), gently mixed and further incubated for 12-18 hours at 37°C. Cells were washed with phosphate buffered saline (PBS; Gibco) and crosslinked with 365 nm wavelength bulbs for 0.5 J/cm^2^ (UV Stratalinker, Agilent set to 5000). Following cross-linking, 1 ml of PBS was added to each plate. Cells were scraped, then pelleted by centrifugation at 500g for 5 minutes at 4°C. The supernatant was decanted, and cell pellets were flash frozen with liquid nitrogen and stored -80°C until further processing.

Cell pellets were resuspended to homogeneity on ice in 2.5x the packed cell volume (PCV) of Buffer A (10mM HEPES pH 7.5, 4 mM MgCl_2_, 10 mM KCl, 340 mM sucrose, 10% glycerol, 0.5 mM DTT, 5 mM 2-mercaptoethanol and 1X protease inhibitor cocktail [1mM ABESF, 0.8 μM aprotinin, 20 μM leupeptin, 15 μM pepstatin A, 40 μM bestatin, 15 μM E-64, 600 μM Benzamidine]). To lyse these cells, an equal of volume of Buffer A with 0.2% (w/v) Triton X-100 was added to the cell suspension and mixed by gentle inversion. Cells were incubated on ice for 12 minutes with occasional gentle inversion, then centrifuged at 1,200g for 5 min at 4°C. The cytosolic extract (supernatant) was saved at -80°C and the nuclei were resuspended in 0.4 ml buffer A. The pellet was aliquoted into ∼5 × 10^6^ nuclei per experimental condition and all samples were resuspended in the same volume (400 µl) of sBA. Nuclei were subjected to DNase digestion (50U, Worthington) for 10 min at 37°C to fragment chromatin. Soluble proteins and nucleic acids were extracted by adding NaCl from 5M stock and EDTA from 0.5M stock to final concentrations of 400 and 10 mM, respectively, followed by incubation with gentle end-over-end rotation for 30 minutes at 4°C. Nuclei were centrifuged at 15,000g for 15 minutes at 4°C to pellet chromatin and release the soluble nuclear extract. The clarified soluble nuclear extract was transferred to a new tube and an equal volume of buffer A was added to lower the salt concentration to 200 mM. Then we set aside 5% and 1% v/v input in separate tubes for RNA and protein analysis, respectively. 100 U/ml RiboLock RNase Inhibitor (ThermoFisher) was added to each sample, which represented the first IP input.

Soluble nuclear extracts were subjected to first immunoprecipitation (under native conditions) with 12CA5-conjugated protein A Dynabeads. For 12CA5 conjugates, 50 µl protein A Dynabeads (ThermoFisher) were prepared for each sample by removing the storage buffer and incubating for 30 minutes at room temperature with 1 ml of 12CA5 medium harvested from 12CA5-producing hybridoma. The beads were washed 3X with 1 ml RIP wash buffer (50 mM K_x_H_y_PO_4_ pH 7.2, 5 mM MgCl_2_, 300 mM KCl, 0.01% (w/v) Tween-20, 1X Protease Inhibitor Cocktail, 5 mM 2-mercaptoethanol) and then resuspended in 50 µl of PBS. 12CA5-conjugated Dynabeads were added to nuclear extracts and incubated at 4°C for 2-4 hours with rotation. After IP, the resin was collected by neodymium magnetic rack and washed 3X with 1ml RIP wash buffer, including 2X tube changes at 4°C.

For the second immunoprecipitation, we added 700 µl denaturing buffer (0.009 % Tween-20 (w/v), 200 mM K_x_H_y_PO_4_ pH 7.95, 7.3 M Urea, 18 mM imidazole, 9% glycerol, 0.9M NaCl) directly to beads, and incubated for 5 minutes at 50°C in a shaking incubator 600 rpm, then collected with a magnetic rack. The supernatant was transferred to 20 µl of Cobalt Dynabeads (prewashed with denaturing buffer), and incubated for 5 minutes at room temperature with rotation. Beads were washed 4X with denaturing buffer, including 2X tube changes followed by one wash with 1X with Proteinase K Buffer (0.25% SDS, 10 mM HEPES, pH 7.5, 250 mM NaCl, 10% glycerol, 0.5% Tween-20, final pH 6.85). Beads were re-suspended in 1X original bead volume Proteinase K Buffer. Proteinase K (Roche) was added to 1 mg/ml final concentration and incubated for 40 minutes at 50°C in a shaking incubator at 600 rpm. After magnetic separation, the elution was transferred to a fresh tube and resin was washed with one bead volume of Proteinase K buffer, both elutions were then combined.

To obtain the de-crosslinked RNA, we added an equal volume Phenol:Chloroform:IAA, pH 8.0, and mixed vigorously for 30 seconds, incubated for 1 minute at room temperature, then centrifuged 15,000g for 2 minutes at 4°C. The aqueous layer was used as input for RNA Clean and Concentrator columns (Zymo), and processed according to the manufacturers protocol (including “In-tube” DNase digest), then the purified RNA was eluted in 30 µl RNase-free water and flash frozen in liquid nitrogen. This recovered RNA was used as input for Illumina Library preparation by using the NEB-Next Ultra Directional RNA-seq Library Preparation Kit for Illumina (NEB), then sequenced by HiSeq (Illumina) at the University of Chicago Functional Genomics Core.

### Nuclear fractionation coupled to RNase treatment

One 10 cm plate of FLAG-WDR5 T-REx™ 293 cells at 70-80% confluency was used per replicate for the nuclear fractionation^128^. Cells were washed twice with PBS, scraped in 1 ml PBS on ice and centrifuged at 500g for 10 minutes at 4°C. Cells were resuspended in 3 packed cell volumes (PCV) of hypotonic buffer HB (10 mM HEPES pH 7.8, 10 mM KCl, 1.5 mM MgCl_2_, 340 mM sucrose, 10% glycerol (v/v), 0.5 mM PMSF, 0.5 mM DTT, and 1X protease inhibitor cocktail). Cells were lysed by adding an equal volume of HB supplemented with 0.2% Triton-X100 (v/v) with incubation at 4°C with gentle rotation for 12 minutes. Nuclei were collected by centrifugation at 500g for 10 minutes at 4°C. The supernatant was saved as the cytoplasmic fraction. The crude nuclei were resuspended in 3 PCV of hypotonic buffer HB and cleaned through a sucrose cushion (5 ml; 30% w/v sucrose, HEPES, pH 7.8, 1.5 mM MgSO_4_) with centrifugation at 1300g for 12 minutes at 4°C. Recovered nuclei were resuspended in 4 PCV of hypotonic buffer HB and divided equally in three tubes. Equal volume of HB supplemented to 300 mM, 500 mM, or 700 mM KCl was added to achieve the final concentration of KCl at 150, 250 and 350 mM, respectively. Each nuclei suspension was further split in two equal volumes for treatment or no treatment with RNase cocktail consisting of RNase A (Thermo, 10µg), RNase I (Thermo, 10U), RNase T1 (Thermo, 1000U) and RNase H (NEB, 5U). Nuclei suspensions were then subjected to gentle rotation at 4°C for 2 hours to isolate soluble nuclear extract. Extracted nuclei were recovered by centrifugation at 8000g for 5 minutes at 4°C. The supernatant was clarified by high-speed centrifugation at 18000g for 30 minutes at 4°C and saved as the nucleoplasm fraction. To solubilize the chromatin pellet, the extracted nuclei were resuspended in 2 PCV of buffer HB supplemented to 1 mM CaCl_2_ pre-equilibrated briefly at 37°C and then subjected to micrococcal nuclease digestion (0.15 U/μL, Worthington) for 25 minutes. This digestion was quenched by adding SDS-PAGE loading buffer. Samples were loaded for SDS-PAGE with equivalent volume, as both the soluble nuclear and chromatin pellet extracts were made from the same nuclei prep for a given salt concentration, and equivalent numbers of nuclei were used for each salt concentration. Gels were transferred to PVDF (Millipore, Immobilon-P SQ) in modified Towbin’s buffer (25 mM Tris pH 8.0, 192 mM glycine, 20% Methanol, 0.04% SDS [w/v]), by semi-dry apparatus (Bio-Rad Trans-Blot SD) at 22V for 23 minutes. Membranes were then blocked for at least one hour in 2% (w/v) Amersham ECL Prime blocking agent (Cytiva) in TBST. Primary antibodies in the same blocking solution were used as follows: 1:1000 α-CDK8 (Abcam 229192), 1:1000 α-FLAG (Sigma M2), 1:1000 α-GAPDH (Cell Signaling 5174), 1:3000 α-H3 C-term (Epicypher 13-0001), 1:1000 α-SF3B155 (Bethyl A300-996A). Secondary goat anti-mouse (1:10,000; Thermo 31432) and goat anti-rabbit (1:10,000; Sigma A6154) HRP-conjugated antibodies were detected using ECL Select (Cytiva; RPN2235) on a Fuji LAS-4000 imager.

### Protein expression and purification

Recombinant tobacco etch virus protease (TEV)-cleavable N-terminal His_6_-tagged human WDR5^79^ was overexpressed in *Escherichia coli* (Rosetta2-DE3-pLysS cells) by growing cells at 37°C in LB medium containing appropriate antibiotic, to an OD_600_ of 0.4-0.6. The temperature was then lowered to 16°C, and cells were induced with 1mM isopropyl-1-thio-D-galactopyranoside (IPTG) for 16-18 hours. Cells were harvested, resuspended in lysis buffer (50 mM sodium phosphate pH 8.0, 500 mM NaCl, 10 mM imidazole, 10% glycerol, 1 mM PMSF, 10 mM 2-mercaptoethanol, 1X protease inhibitor cocktail) and lysed by passing through an Emulsiflex C3 homogenizer, and clarified by centrifugation. The clarified lysate was incubated with Ni-NTA agarose slurry (Qiagen, 1 mL slurry per liter of culture) in a 2.5 × 10 cm Econo-Column (Bio-Rad) for 30-45 minutes at 4°C, and was washed by the sequence of 10 ml of high salt wash buffer (0.9X lysis buffer with 1M NaCl), 10 ml lysis buffer, 10 ml low salt wash buffer (lysis buffer with 250 mM NaCl), followed by 3 × 5 ml of elution buffer (0.9X low salt buffer with 300 mM imidazole). The elutions were pooled together and diluted with 3 volumes of 150 mM Tris•HCl (pH 7.0), and loaded on a heparin HiTrap column (5mL, GE Healthcare) that was equilibrated with the buffer A (50 mM Tris•HCl pH 7.0, 0.1 M NaCl, 5 mM 2-mercaptoethanol). The protein was eluted using a gradient of buffer B (50 mM Tris•HCl pH 7.0, 1M NaCl, 5 mM 2-mercaptoethanol) over 10 c.v. Eluted protein was treated with TEV protease^129^(1:100 w/w TEV:WDR5) at 4°C overnight in the presence of 3 mM DTT and 2 mM EDTA and analyzed on an SDS-PAGE gel to confirm hexahistidine cleavage. Finally, the pooled fractions were purified on an Superdex 75 column (10/300 GL, Cytiva) into the final filter-binding buffer (50 mM Tris•HCl pH =7.0, 150 mM NaCl, 5 mM MgCl_2_, 0.1 mM EDTA, 10 mM 2-mercaptoethanol). All WDR5 mutants were also purified using the same procedure.

Human MLL1 core complex components were individually expressed and purified as follows. N-terminal SUMO-tagged RBBP5 containing pET28a plasmid was a generous gift from the Dou lab^130^. N-terminal GST-MLL1 (3745-3969), His_6_-ASH2L, and His_6_-DPY30 were generous gifts from the Cosgrove lab^25,89^. All proteins were expressed similarly to WDR5 except RBBP5 and Ash2L, which were induced at 0.1 mM IPTG and 0.25 mM IPTG, respectively. For RbBP5, post-Ni-NTA column elution, the pooled fractions were treated with ULP1 (a generous gift from Staley Lab, University of Chicago) at 4°C overnight in the presence of 3 mM DTT and 2mM EDTA. The cleaved protein was purified using MonoQ (1mL, GE Healthcare) or HiTrapQ chromatography (5 mL, GE Healthcare). For ASH2L, the pooled Ni-NTA elutions were purified on a Q fast-flow column (5 mL, GE Healthcare) followed by overnight TEV cleavage. GST-tagged MLL1 lysate was loaded on a Hi-Trap GST column (5mL, GE Healthcare), washed with 10 c.v. high salt wash buffer (50 mM Tris•HCl pH 7.0, 500 mM NaCl), followed by overnight on-column TEV cleavage (300 µL of 1mg/mL TEV for 1L MLL1 prep), and elution in binding buffer (50 mM Tris•HCl pH 7.0, 500 mM NaCl, 10 mM 2-mercaptoethanol). The eluted protein was diluted with 3 volumes of buffer A (50 mM Tris•HCl pH 7.0) and further purified over a Heparin HiTrap (5mL, GE Healthcare) column to get rid of bound nucleic acids. DPY30 was purified over a Ni-NTA column, followed by overnight TEV cleavage. All proteins were additionally purified over an Superdex 75 size exclusion column into the following buffer: 20 mM Tris pH 7.5, 300 mM NaCl, 1 μM ZnCl_2_. The final fractions were collected and concentrated to ∼2mg/mL for each protein, and flash frozen in 20 mM Tris pH=7.5, 300 mM NaCl, 1 μM ZnCl_2_ in 40-50% glycerol and stored at -80 °C until use.

### *In vitro* transcription and radiolabeling

DNA templates for different RNAs were generated using PCR with a T7 promoter sequence (TAATACGACTCACTATTA) in the forward primer. The DNA templates products were purified using DNA Clean and Concentrator kit (Zymo Research) and directly used for *in vitro* transcription. Each body labeled transcription reaction (20 μL) contained 1 μg digested or PCR purified DNA, 3 mM A/C/GTP, 0.3 mM UTP, 2.5 μL ^32^P-labeled ⍺-UTP (Perkin Elmer BLU007H250UC), 200 mM HEPES (pH 8.0), 12.5 mM MgCl_2_, 1 mM DTT, 2 mM spermidine, and 10 µg T7 RNA polymerase (P266L mutant, plasmid a kind gift of Nicholas Reiter). The reaction was allowed to proceed at 37°C for 2 hours, followed by quenching with the addition of 5 units Turbo-DNase (Invitrogen) for 15 minutes, followed by addition of EDTA to 25 mM. The RNA was purified using the RNA Clean and Concentrator kit (Zymo Research), eluted in RNAse-free water and stored at -20°C until use. Transcriptions of constructs shorter than 400 nucleotides were quenched by using 2X formamide loading buffer (95% formamide and 25 mM EDTA, 0.03% (w/v) xylene cyanol and bromophenol blue), and were subsequently purified on 5 or 8% polyacrylamide midi gel (Gibco V16). The RNA band was visualized using UV shadowing, and the gel slices were incubated overnight in TEN_250_ buffer (10 mM Tris•HCl pH=8.0, 1 mM EDTA, 250 mM NaCl). The RNA was precipitated using 3 volumes of ethanol, and centrifuged at 16000g and the pellet was washed with 70% ethanol, followed by air-drying. The RNA was resuspended in RNase-free water, and stored at -20°C until further use.

### Filter-binding assays

All the membranes were equilibrated in the filter binding buffer (50 mM Tris•HCl pH 7.0, 150 mM NaCl, 5 mM MgCl_2_, 0.1 mM EDTA, 10 mM 2-mercaptoethanol) for >30 minutes. Reactions were assembled in a 100 μL final volume. Independent serial dilutions for different concentrations of purified protein were prepared in the filter binding buffer and incubated on ice until the start of RNA denaturation. For competitive filter binding assays with the monobodies, increasing concentration of monobody was titrated to a mixture containing filter binding buffer and a constant concentration of WDR5 protein indicated (150 nM, 250 nM or 500 nM). For folding of RNA, 450 μL of filter binding buffer (without MgCl_2_) was added to 1 μL of radiolabeled RNA (>20,000 cpm/μL) and denatured at 90°C for 2 minutes. The RNA was allowed to slowly cool to room temperature for 5 minutes, followed by addition of MgCl_2_ (5 mM), and incubation at 37°C for 30 minutes. 25 μL of the incubated RNA (<< 1 nM) was added to each reaction containing protein and buffer mix. The RNA-protein mix was allowed to incubate at room temperature for 30 minutes, following which the reactions were resolved by filtration through pre-washed (with filter binding buffer) stack of nitrocellulose (Bio-Rad), and Hybond-N^+^ (GE Healthcare) or Zeta-Probe (Bio-rad) membranes backed by Whatman paper, using a S&S Minifold-1 dot blot system (Whatman Schleicher & Schuell). After filtration of binding solutions, each well was washed twice with 100 μL washing buffer (filter binding buffer without MgCl_2_). The membranes were air-dried, exposed to a phosphor screen through saran wrap, and scanned on a Typhoon 9200 (GE Healthcare Life Sciences). Data were quantified by TotalQuant with rolling ball 150 background correction and normalized to min-max lane values. Equilibrium dissociation constants, *K*_d_, were obtained by nonlinear regression of the binding data fitted to a one-site binding model [Y=B_max_*X^n/(*K*_d_^n + X^n)]. Computed *K_d_* and n for each RNA are provided in Table S1.

For RNA-RNA competition assays, [^32^P] body-labeled RNA was incubated with excess WDR5 (∼4-fold higher concentration than the *K*_d,app_ of the respective labeled RNA), along with the indicated titrations of competing unlabeled “cold” RNA for 30 minutes at room temperature. The samples were processed as above after the incubation.

### Fluorescence polarization (FP) assay

For protein-RNA FP assay, purified proteins were dialyzed into the binding buffer (50 mM Tris•HCl pH 7.0, 150 mM NaCl, 5 mM MgCl_2_, 0.1 mM EDTA, 10 mM 2-mercaptoethanol) for 12 hours with one change of dialysis buffer after 4 hours. Indicated serial dilutions of protein were incubated with 10-30 nM 5’-FAM labeled RNA oligonucleotide, suspended in the binding buffer. Independent dilutions were made for replicated measurements. The dilutions were incubated at room temperature for 30 minutes as longer incubations did not apparent binding. FP was measured at room temperature on a TECAN Infinite 200 Pro plate reader with excitation/emission wavelengths of 485 (20) nm/535 (25) nm. Readings were corrected for G-factor calculated from an RNA only sample with binding buffer set as the blank. The reference polarization of the RNA only sample was set to 20 mP. Normalized polarization were regressed with the in-built GraphPad Prism equation for “Specific binding with Hill slope” [Y=B_max_*X^n/(*K*_d_^n + X^n)] to calculate *K*_d, app_ for the interaction. Computed *K_d_* and n for each RNA are provided in Table S1.

For testing the competition between RNA species using FP, the indicated highest concentration of the unlabeled competitor RNA oligonucleotide was added to a pre-reaction mixture of WDR5 (4 µM) and 5’-FAM labeled RNA (10 nM). This mixture was serially diluted with the diluent mixture consisting of WDR5 (4 µM) and 10 nM labeled RNA to obtain the desired concentration range of the competitor RNA. Reactions were incubated at room temperature for 30 minutes and FP readings were taken as described above.

For FP of WDR5 with the MLL1 peptide, 5’-FITC labeled MLL1 peptide (FITC-HGSARAEVHLY) was synthesized by the Tufts University Core Facility. The peptide was dissolved in 30% acetic acid to make a stock solution of 0.5 mM peptide. FP binding assays were performed with 10 nM peptide diluted in the binding buffer and the indicated concentration range of WDR5. FP competition assays were done by serially diluting the RNA with a pre-reaction mixture of 10 nM MLL1 peptide and 250 nM WDR5. Reactions were incubated for 30 minutes at room temperature. Data were collected and processed as described above.

### Simulation of competitive binding in fluorescence polarization

The competitive binding between RNA of different lengths was simulated using the script from https://github.com/stephenfloor/competitive-binding-simulator based on equations from Roehrl et al^131^. The values used were -kd 7 -rt 4 -lt 0.01 -ki 0.58 (for 500 nt), 0.8 (for 250 nt) and 3 (for 50 nt).

### *In vitro* histone methyl transferase assays

Each purified component MLL1_3745-3969_, WDR5, RBBP5, ASH2L, and DPY30, were mixed in 1:1:1:1:2 stoichiometry with KMT assay buffer (50 mM Tris•HCl pH 8.0, 50 mM NaCl, 2.5 mM MgCl_2_, 1 mM DTT), and allowed to incubate on ice for 30 minutes prior to start of the experiment. The RNA was heated at 90°C for 2 minutes followed by cooling at room temperature for 5 minutes.

0.5 μM stock of RNA was made in 1X KMT buffer and the RNA was renatured at 30°C for 10 minutes. Each 20 μL reaction consisted of 1 μM final concentration of MLL1 complex, 1X KMT assay buffer, 5 μM H3 peptide (Sequence: H3 1-20, plus C-terminal tyrosine for UV quantification),1 μM Adenosyl-L-Methionine, S-[methyl-^3^H] (Perkin Elmer NET155H250UC), and indicated concentration of RNA. The appropriate volumes of MLL1 complex and 1X KMT buffer were added to each reaction mixture and incubated at room temperature for 5 minutes, followed by addition of RNA. The MLL complex-RNA mix were allowed to incubate at room temperature for 30 minutes, and the methyltransferase reaction was initiated by adding the MLL1-RNA mix to a pre-mixed solution of 5 μM unmodified H3 (1-20+Y peptide) substrate and 1 μM ^3^H-SAM (Perkin Elmer). 4 μL of reaction was taken indicated intervals, and the reaction was quenched by immediately spotting the reaction mixture on P81 phosphocellulose paper. The P81 paper was allowed to air-dry, washed four times with freshly prepared 50 mM NaHCO_3_, and ^3^H incorporation measured by using Liquid Scintillation Counter.

### Circular dichroism

The protein was diluted to 3 μM in 10 mM sodium phosphate buffer (pH 7.1). Circular dichroism spectra were recorded on a Jasco J-715 CD Spectrometer equipped with a thermoelectrically controlled cell holder using a quartz cell with a 1.0 mm optical path length. For the melt curve analysis, the proteins were continuously scanned over a temperature range from 20°C to 90°C with an increment of 0.5°C.

### Differential scanning fluorimetry

For fluorescence-based thermal shift assays^132^, proteins were diluted to 1 μM in TS buffer (50 mM NaCl, 50 mM Tris pH =8.0, 2.5 mM MgCl_2_) and 2X Sypro Orange (ThermoFisher). The temperature range of the thermal melt was 30°C to 90°C, with 0.5°C increment and a 10 second stay at each temperature. The fluorescence was measured on a Bio-Rad CFX96 plate reader in FRET mode. The melting temperature was calculated from the inflection point of the linear range of fluorescence increase plot (also corresponds to the local maxima of the first-derivative of the fluorescence vs temperature plot).

### Generation of monobodies

The procedure of selecting target specific monobodies from phage and yeast display libraries has been described previously^91^. The first selection cycle resulted in Mb(WDR5_S4) and other clones that target MLL-interaction site on WDR5^92^. To identify monobodies that bind to a distinct epitope, the yeast libraries were rescreened in the presence of 100 nM Mb(WDR5_S4) as saturating competitor by fluorescence-activated cell sorting. A binding assay for testing the affinity of the individual monobody clones was performed using yeast surface display as described previously^133^.

### Isothermal Calorimetry

Isothermal titration calorimetry (ITC) experiments were performed using MicroCal iTC200 calorimeter (MicroCal), at 25°C in a sample buffer (25 mM HEPES pH 8, 50 mM NaCl). ML17 monobody (75 µM) solution was injected into WDR5 (8.3 µM) solution in the sample cell, while being stirred at 750 rpm. A 60 second delay at the start of the experiment was followed by 20 injections of 2 μl each of the titrant solution, spaced 180 second apart. Blank injections of the ML17 monobody into buffer were subtracted from the experimental titrations, and binding isotherms were fit to a theoretical one binding site per titrant model. A nonlinear best-fit binding isotherm for the data was used to calculate the dissociation constant using the Origin 7.0 software (Origin-Lab Corp.).

### GST-pulldown assay

Pierce Glutathione Magnetic Agarose Beads (ThermoFisher; 15 µL per reaction) were washed twice with the binding buffer (50 mM Tris pH 8.0, 150 mM NaCl, 5 mM MgCl_2_, 0.1 mM EDTA, 10 µM ZnSO_4_, 10% Glycerol, 0.01% NP-40, 5 mM 2-mercaptoethanol, 2mM DTT, 1mM PMSF) and blocked overnight with 1 mg/ml BSA. Blocked beads were washed once with the binding buffer. 250 pmol of purified GST-MLL1 was added to these beads in the binding buffer and incubated at 4°C for 1 hour with rotation. In parallel, 75 pmol of WDR5 was incubated with the indicated amount of RNA in 45 µl reaction volume at 4°C for 1 hour with rotation. GST-MLL1 bound beads were washed twice with the binding buffer. WDR5-RNA mixture was then added to beads and incubated at 4°C for 1 hour with rotation. Unbound material was saved from the supernatant and beads were washed three times with the binding buffer with tube changes. The bound material was eluted by either heating the beads in 20 µl of 1X SDS loading dye or incubation with 20 µl of the binding buffer supplemented with 50 mM glutathione. Protein samples were run on SDS polyacrylamide gels followed by staining with SYPRO-Ruby as per the product instructions for the high sensitivity overnight staining protocol. Gels were imaged on Typhoon 5 (Cytiva) with excitation/emission at 488 nm/670 (30) nm and quantified in ImageJ.

### *In situ* RNA competition assay

One 10 cm plate of FLAG-WDR5 T-REx™ 293 cells at 70-80% confluency was used per replicate for the nuclear fractionation^128^. Cells were washed twice with PBS, scraped in 1 ml PBS on ice and centrifuged at 500g for 10 minutes at 4°C. Cells were resuspended in 3 packed cell volumes (PCV) of hypotonic buffer HB (10 mM HEPES pH 7.8, 10 mM KCl, 1.5 mM MgCl_2_, 340 mM sucrose, 10% glycerol (v/v), 0.5 mM PMSF, 0.5 mM DTT, and 1X protease inhibitor cocktail). Cells were lysed by adding an equal volume of HB supplemented with 0.2% Triton-X100 (v/v) with incubation at 4°C with gentle rotation for 12 minutes. Nuclei were collected by centrifugation at 500g for 10 minutes at 4°C. The supernatant was saved as the cytoplasmic fraction. The crude nuclei were resuspended in 3 PCV of hypotonic buffer HB and cleaned through a sucrose cushion (5 ml; 30% w/v sucrose, HEPES, pH 7.8, 1.5 mM MgSO_4_) with centrifugation at 1300g for 12 minutes at 4°C. Recovered nuclei were resuspended in 6 PCV of hypotonic buffer HB supplemented to 150 mM KCl and divided equally in three tubes. In vitro transcribed RNA was added to a final concentration of 1 µM. Nuclei suspensions were then subjected to gentle rotation at 4°C for 1.5 hours to isolate soluble nuclear extract. Extracted nuclei were recovered by centrifugation at 8000g for 5 minutes at 4°C. The supernatant was clarified by high-speed centrifugation at 18000g for 30 minutes at 4°C and saved as the nucleoplasm fraction. To solubilize the chromatin pellet, the extracted nuclei were resuspended in 2 PCV of buffer HB supplemented to 1 mM CaCl_2_ pre-equilibrated briefly at 37°C and then subjected to micrococcal nuclease digestion (0.15 U/μL, Worthington) for 25 minutes. This digestion was quenched by adding SDS-PAGE loading buffer. Samples were loaded for SDS-PAGE with equivalent volume, as both the soluble nuclear and chromatin pellet extracts were made from the same nuclei prep for a given RNA treatment, and equivalent numbers of nuclei were used for each RNA treatment. Gels were transferred to PVDF (Millipore, Immobilon-P SQ) in modified Towbin’s buffer (25 mM Tris pH 8.0, 192 mM glycine, 10% Methanol, 0.04% SDS [w/v]). Membranes were then blocked for at least one hour in 2% (w/v) Amersham ECL Prime blocking agent (Cytiva) in TBST. Primary antibodies in the same blocking solution were used as follows: 1:1000 α-ASH2L (Bethyl A300-107A), 1:1000 α-FLAG (Sigma M2), 1:1000 α-GAPDH (Cell Signaling 5174), 1:20,000 α-H3 C-term (Epicypher 13-0001), 1:1000 α-MLL1-C (Cell Signaling 14197), 1:2000 α-Pol II S5P (Cosmo Bio, MCA-MABI0603-100-EX), and 1:1000 α-RBBP5 (Bethyl, A300-109A). Secondary goat anti-mouse (1:10,000; Thermo 31432) and goat anti-rabbit (1:10,000; Sigma A6154) HRP-conjugated antibodies were detected using ECL Select (Cytiva; RPN2235) on a Fuji LAS-4000 imager.

## QUANTIFICATION AND STATISTICAL ANALYSIS

### Computational analyses

For PAR-CRAC, reads were aligned to the hg38 genome using hisat2 aligner^134^ with --score-min L,0,-0.8 option. Duplicated reads were removed using the MarkDuplicates function of Picard Toolkit^135^ with REMOVE_DUPLICATES=true option. The deduplicated bam files were converted to counts per million (CPM)-normalized bigwig and bedgraph files by taking only primary alignments of all mapped reads into account, using the bamCoverage function of deepTools (−bs 1 --samFlagExclude 256 --normalizeUsing CPM) ^136^. The bedgraph files were then used to score the PAR-CRAC counts on the defined genomic regions (such as isoform consolidated genes) using bedtools (map -a consolidated_gene.bed -b CPMnormlized.bg -c 4 -o sum). Total RNA-seq dataset from HEK293 (SRR650317) cells was processed similarly and used for input counts. Input normalized PAR-CRAC counts were then used to rank genes for WDR5 PAR-CRAC enrichment. For re-analysis of WDR5 denaturing CLAP-seq^62^, fastq reads were obtained from the GEO database (WDR5 CLAP rep1: SRR27597141, WDR5 CLAP rep2: SRR27597140, Input: SRR27597179) and processed similarly to PAR-CRAC analysis as above.

To quantitate T>C conversion in PAR-CRAC dataset, reads were aligned to the hg38 genome using Bowtie1 (recommended for PARalyzer^68^) with -v 2 -m 10 --best --strata options. The aligned bam files were then used to quantify T>C conversion against hg38 genome using PARalyzer^68^ with minimum count of 1. From the PARalyzer output, genomic regions were scored for the ‘ReadCount’ metric into a bedgraph file. The bedgraph files were converted to bigwig files using bedGraphToBigWig package^137^.

To search for any enriched RNA motif in the WDR5 PAR-CRAC dataset, findMotifs.pl module of the HOMER suite^138^ was used with default options with clusters of 46 nt sequences from PARalyzer with minimum readcount conversion of 2 as sample and fasta file of hg38 sequence as background.

Gene biotypes of top 5% of genes, ranked on the basis of WDR5 PAR-CRAC enrichment (signal/input), were obtained using the ensembldb v2.28.0 and AnnotationHub v3.0 packages in R. Genes falling in ‘transcribed_processed_pseudogene’, ‘transcribed_unitary_pseudogene’, ‘transcribed_unprocessed_pseudogene’, ‘unprocessed_pseudogene’, ‘pseudo’, ‘processed_pseudogene’ categories were merged into ‘pseudo gene’ category for simplifying the representation.

Sites of RNA G-quads (rG4) were obtained from rG4-seq analysis^139^ (GSE77282) and lifted from hg19 to hg38 using the UCSC genome browser.

WDR5 ChIP peaks were obtained by liftover of peaks called in hg19^97^ to hg38 using the UCSC genome browser. For quantitating WDR5 ChIP signal at defined genomic regions, Fastq files for WDR5 ChIP-seq in HEK293 cells^97^, downloaded from the GEO database (GSE60897), were aligned to the hg38 genome using bowtie2 with default options. The bam files were converted to CPM normalized bedgraph files which were used to count WDR5 ChIP signal at upstream regions (−1000 bp relative to the CDS) of the isoform consolidated genes.

Fastq files for our previously generated H3K4me3 ICeChIP in HEK293 cells^140^ were aligned to the hg38 genome using bowtie2 with --sensitive option.

Analysis of chromatin- and nucleoplasm-enriched RNA in HEK293 cells was done as described previously^69^. The intersection of WDR5 ChIP peaks and chromatin-enriched RNA in HEK293 cells was done using bedtools intersect with default options.

Fastq files for MLL1 ChIP-seq in HEK293 cells^141^ were downloaded from the GEO database (Input: SRR4880403, MLL1 IP: SRR4880407) and aligned to hg38 genome using bowtie2 with default options. Input normalized bedgrpah file was generated using bamCompare function of deeptools (-bs 1 --normalizeUsing CPM -b1 IP.bam -b2 INPUT.bam -o output.bedgraph -of bedgraph --pseudocount 1 --operation ratio --scaleFactorsMethod None).

The metagene gene plots were created using the CPM-normalized bigwig files (binned at 1 base pair) of aligned reads to score mean coverage over the region of interest using the computeMatrix function of deepTools in reference-point mode (--missingDataAsZero) and plotted using plotProfile function.

## KEY RESOURCES TABLE

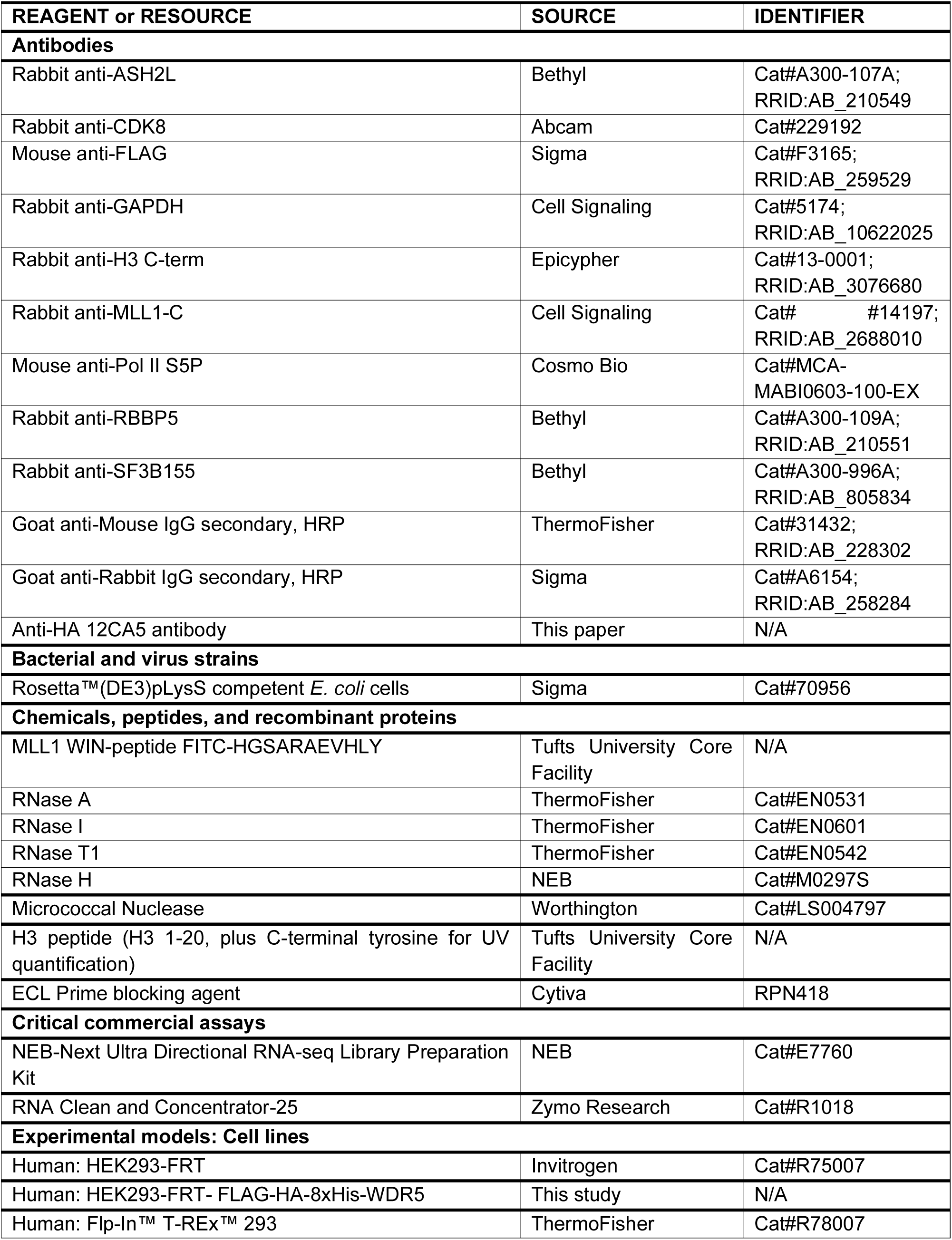
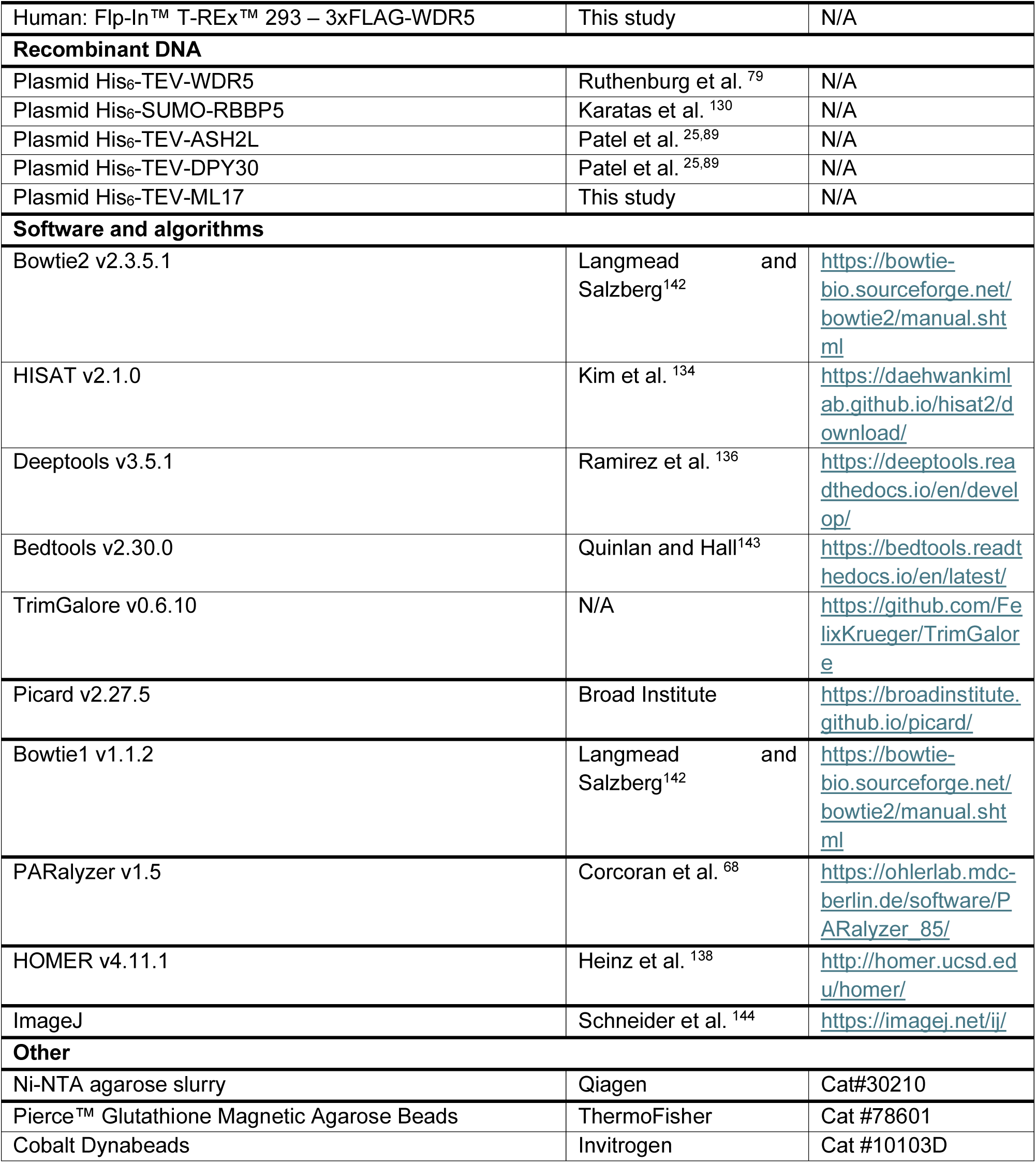

