## Supplemental Document for "Promiscuous RNA binding by WDR5 remodels the KMT2A (MLL1) histone methyltransferase complex to an inactive state"

#### SUPPLEMENTARY FIGURE LEGENDS

**Figure S1. Analysis of cellular RNA-binding to WDR5, related to Figure 1**
**(A)** Western blot analysis of WDR5 in the input, native $1^{st}$ IP, resin and $2^{nd}$ IP of WDR5 PAR-CRAC protocol.
**(B)** Scatter plot of WDR5 PAR-CRAC versus WDR5 denaturing CLAP signal in HEK293 cells. Only the genes with non-zero count in both datasets are shown. Colors correspond to kernel density estimations of scatter plot distribution. r, Spearman correlation coefficient.
**(C)** Proportion of different classes of RNA bound to WDR5. RNA within top 5% of the WDR5 PAR-CRAC enrichment with respect to the input are classified here.
**(D)** Scatter plot of raw counts versus T > C counts in WDR5 PAR-CRAC data. Colors correspond to kernel density estimations of scatter plot distribution.
**(E)** The most abundant motif enriched in the WDR5-bound RNA in PAR-CRAC accounts for only 3% sites.
**(F)** Average count of WDR5 PAR-CRAC thymidine to deoxycytidine (T > C) count centered around RNA G-quads (rG4) or shuffled sites in the genome. The rG4 sites are obtained from Kwok et al. [1] and shuffled sites represents average of five shuffles.
**(G)** Scatter plot of WDR5 denaturing CLAP signal versus the indicated parameter of genes. CPM, counts per million mapped reads; CPKM, counts per thousand base pair of gene per million mapped reads. Only the genes with non-zero count in the comparison dataset are shown. Colors correspond to kernel density estimations of scatter plot distribution. r, Spearman correlation coefficient.
**(H)** Scatter plot of WDR5 PAR-CRAC versus RNA count in the chromatin fraction (*left*) and the nucleoplasm fraction (*right*) from Werner et al. [2] colored according to the length of genes.
**(I)** Rank ordered distribution of genes (with non-zero count) arranged in the ascending order of WDR5 PAR-CRAC enrichment with respect to input.
**(J)** Average count of RNA from CPE and SNE fraction in three replicates around WDR5 ChIP peaks (± 1kb) and equivalent sized shuffled regions of genome. The counts for shuffled region represent average of five shuffles.
**(K)** Representative blot showing the presence of indicated proteins in the cytoplasm (cyto), nucleoplasm and chromatin fractions after nuclear extraction with the indicated salt concentration in the absence or presence of RNase (A, I, T, H).

### Supp Figure 1

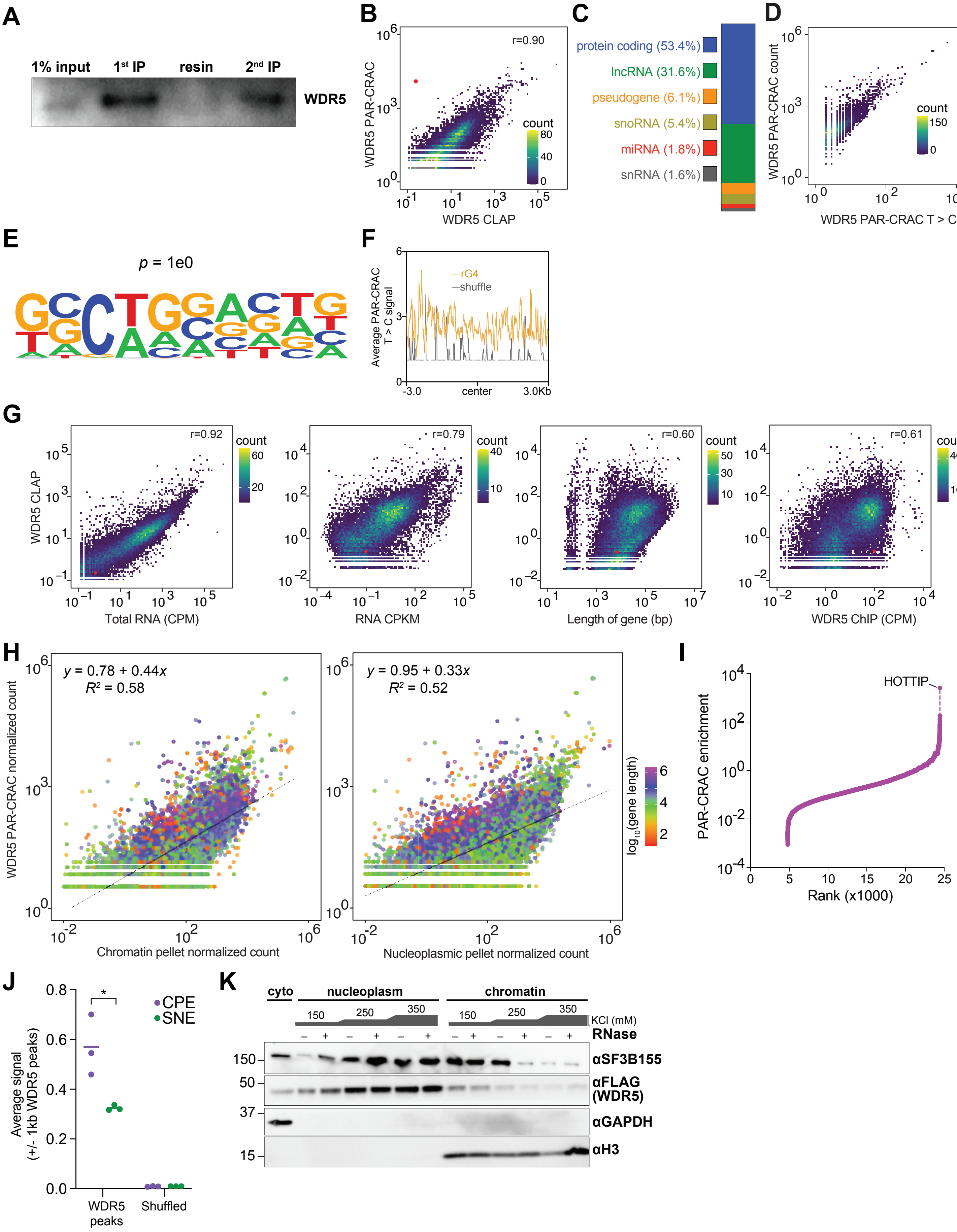

**Figure S2.** ***In vitro*** **WDR5 binding of RNA and supporting experiments, related to Figure 2**
**(A)** Histone methyl transferase assays measuring the transfer of Adenosyl-L methionine, S-[methyl-$^{3}H$] to H3 (1-20) peptide substrate using liquid scintillation counting of phosphocellulose-captured and washed material shows that the reconstituted MLL1 (1 μM) complex is enzymatically active.
**(B)** Equilibrium filter binding measurements of only WDR5 or complete MLL1 KMTase core complex with full length HOTTIP RNA.
**(C)** WDR5 binding curves for different species of indicated RNA. Fluorescence polarization assay was used for ARID1A and $HOTTIP_{24}$; equilibrium filter binding measurements were used for all other RNAs. See Figure 2A for details about these run-off transcribed RNA.
**(D)** Electrophoretic Mobility Shift Assay (*left*) and filter binding dot blot assay (*right,* nitrocellulose and HyBond zeta, top and bottom panel, respectively) of WDR5 with a 24 nt fragment of HOTTIP schematically depicted in Figure 2A.
**(E)** Prediction of $K_d$ of indicated RNAs of different lengths from Monte Carlo simulations.
**(F)** Filter binding membranes (nitrocellulose and HyBond zeta, top and bottom panel, respectively) showing that $HOTTIP_{E1\text{-}3}$ and $HOTTIP_{E1\text{-}2}$, are able to competitively remove $HOTTIP_{E1\text{-}3}$, $HOTTIP_{E1\text{-}2}$, and $HOTTIP_{E3}$ bound to WDR5, indicating that these RNA bind WDR5 with overlapping interfaces. A $HOTTIP_{24}$, a 24 nt RNA that weakly binds WDR5 (Figure S2D) is unable to effectively compete at concentration as high as 500 nM. The concentration of WDR5 is ~4-fold > $K_{d,app}$ for $HOTTIP_{full}$-WDR5 interaction. The concentration of competing RNA is 220 nM, 110 nM, 55 nM and 0 nM for $HOTTIP_{E1\text{-}3}$ and $HOTTIP_{E1\text{-}2}$, and 500 nM, 250 nM, 125 nM, and 0 nM for $HOTTIP_{E3\text{-}24}$ RNA.
**(G)** *In silico* simulations of competitive binding of the indicated competitor to WDR5 (4 μm) displacing the 30 nt ARID1A RNA (10 nM).
**(H)** Comparison of WDR5 affinity to the pertinent binding partners. MLL1-4 and SET1A-B data from Zhang et al. [3], H3K4me2 from Ruthenburg et al. [4] and RBBP5 data from Börgel et al. [5].

### Supp Figure 2

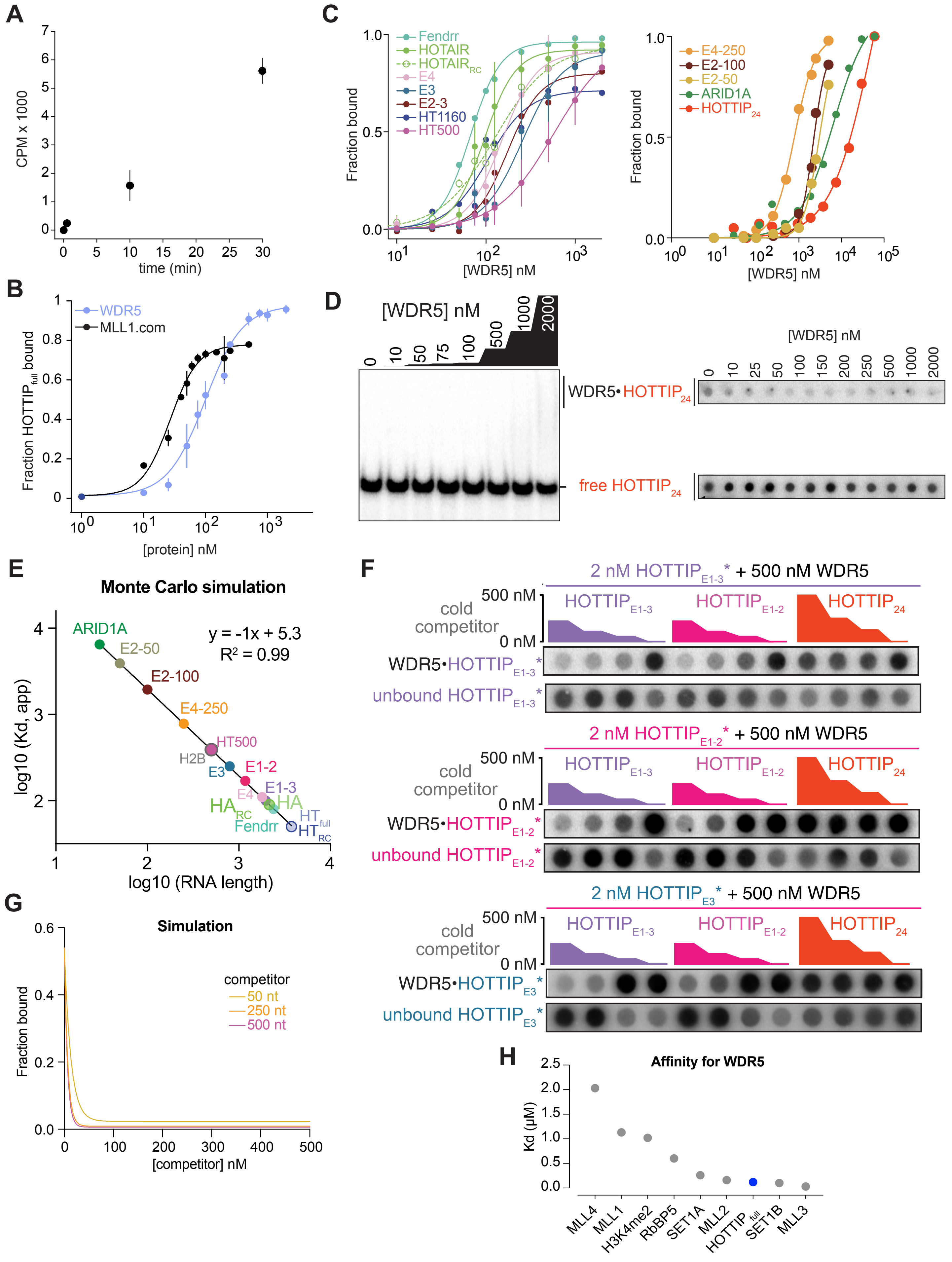

**Figure S3. Probing the RNA-binding interface of WDR5 , related to Figure 3**

**(A)** Derivative plots of thermal optical melting profiles using circular dichroism of WDR5 and its mutants (F266A, Side Face, and TSF (top face + side face)).

**(B)** Structure of WDR5 (PDB ID: 4ESG) indicating the mutated residue sites.

**(C)** Equilibrium filter binding measurements showing complete binding curves for the indicated mutants of WDR5 to full length HOTTIP RNA.

**(D)** Affinity of the indicated WDR5 mutants obtained from equilibrium filter binding measurements to full length HOTTIP RNA presented at relative $K_d$ to a side-by-side contemporaneous WT WDR5 preparation. See panel B and C for the nomenclature and location of the mutations. Note that K250 is involved in RBBP5 [6] binding and was previously noted to impact RNA binding [7].

**(E)** Structure of WDR5 indicating the interacting surface of ARID1A RNA (low confidence AlphaFold3 prediction). WDR5 residues within 4.5 Å of any atom of the ARID1A RNA are highlighted in green.

**(F)** Structure of WDR5 indicating the interacting surface of $MLL1_{3745\text{-}3969}$ peptide (PDB ID: 7UD5 [8]) WDR5 residues within 4.5 Å distance of any atom of the $MLL1_{3745\text{-}3969}$ peptide are highlighted in green.

**(G)** Histone methyl transferase assays measuring the transfer of $^3$H-labeled S-adenosyl methionine to histone substrate ($H3_{1\text{-}20}$ peptide) using liquid scintillation counting in the presence of indicated concentration of $HOTTIP_{full}$ RNA with MWRAD complex (MLL-SET, WDR5, RBPP5, ASH2L, DPY30).

**(H)** Same as F, except MTRAD (TSF mutant of WDR5) or MRAD (missing WDR5) complex was used.

**(I)** *Left*, Structure of WDR5 indicating the interacting surface of S4 monobody (PDB ID: 6BYN [9]); WDR5 residues within 4.5 Å distance of any atom of the S4 monobody are highlighted in green. *Right*, $HOTTIP_{full}$ RNA bound by WDR5 in the absence or presence of S4 monobody by GST-pull down.

**(J)** Equilibrium filter binding measurements of ML17 monobody to $HOTTIP_{full}$ RNA, indicating no detectable binding up to low µM range.

**(K)** Equilibrium filter binding competitive assays with 500 nM WDR5 in the presence of different HOTTIP RNA fragments with increasing concentration of the monobody. The $IC_{50}$ is calculated to be 2200 ± 1000 nM, 1500 ± 700 nM and 2500 ± 700, for $HOTTIPE_{1\text{-}2}$, $HOTTIP_{E3}$, and $HOTTIP_{500}$ RNA, respectively. The high error and lower $IC_{50}$ than HOTTIP RNA for these RNAs may be attributed to the sub-saturating concentration of WDR5. Shown are average from two replicates with the range between the two as the error. The error in the values of the IC50 is derived from the fit of the curve to the following equation: $f_{bound} = F_{max} - [(F_{max} - F_{min})/(1+[M]/IC_{50})]$, where $f_{bound}$ is the observed fraction of RNA bound to WDR5 at different monobody concentration ([M]), $F_{max}$ is the RNA bound at 0 [M].

### Supp Figure 3

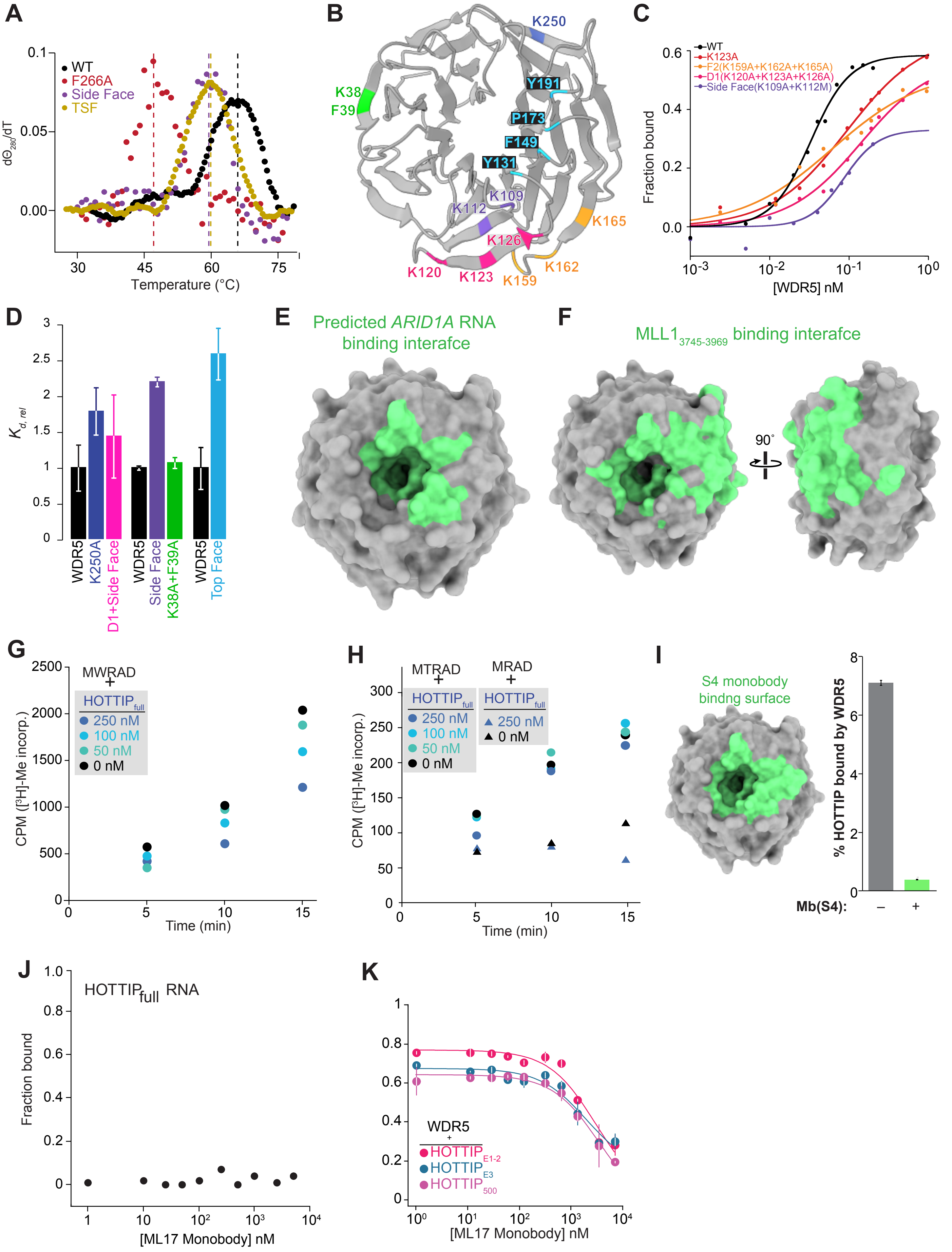

**Figure S4. RNA disrupts MLL1 complex assembly, related to Figure 4**
**Supplement Figure 4. RNA disrupts MLL1 complex assembly**
**(A)** SYPRO-Ruby staining of unbound WDR5 in the supernatant and the $A_{260}$ measurements of the supernatant reflecting unbound RNA/nucleotide in the GST-MLL1 pulldown assay in the presence of the indicated concentration of HOTTIP full length RNA ($HOTTIP_{full}$, *left*) or free NTPs, concentration-matched per-nucleotide to HOTTIP (*middle*, 3754x the indicated concentration to match HOTTIP nucleotide content). *Right*, Quantitation of unbound WDR5 (in supernatant) across the indicated range of RNA/NTP titrations. Shown are the average from two replicates ± S.D.
**(B)** Experimental replicate of assay in Figure 4A. SYPRO-Ruby staining of GST-MLL1 and WDR5 proteins eluted from glutathione resin after GST-MLL1 and WDR5 incubation in the presence of the indicated concentration of HOTTIP full length RNA (HT-FL, *top*) or concentration-matched free NTPs (*bottom*).
**(C)** SYPRO-Ruby staining of GST-MLL1 and WDR5 proteins eluted from glutathione resin after GST-MLL1 and WDR5 incubation in the presence of the indicated concentration of the non-binding 24 nt fragment of HOTTIP.
**(D)** Fluorescence polarization of fluorochrome labeled MLL1 peptide titrated with WDR5.
**(E)** Filter binding membranes (nitrocellulose and HyBond zeta, top and bottom panel, respectively) MM401 is unable to effectively compete with $WDR5\text{-}HOTIP_{full}$.
**(F)** Structure of WDR5 indicating the footprint of MM401 (PDB ID: 4GM9 [10]) and $MLL1_{WIN}$ (PDB ID: 4ESG [11]). The buried surface area between WDR5 and the interacting partner is shown in parentheses.
**(G)** Experimental replicate of assay in Figure 4H. Representative western blot showing the presence of indicated proteins in the cytoplasm (cyto), nucleoplasm and chromatin fraction after fractionation with no RNA or a 500 nt fragment of HOTTIP RNA or a 24 nt fragment of RNA.

#### Supp Figure 4

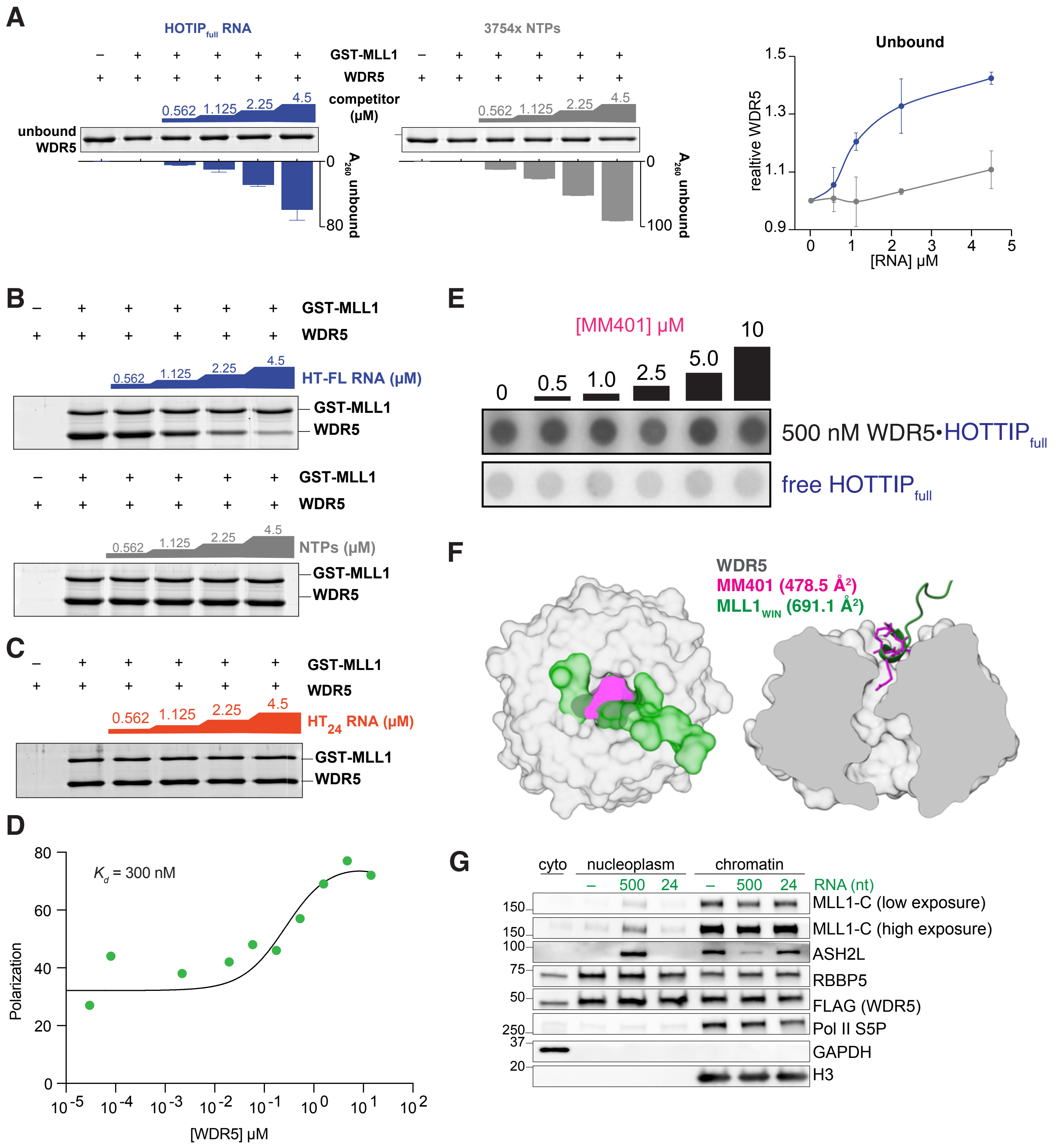
